# *Saccharomyces cerevisiae* Dmo2p is involved in the mitochondrial copper metabolism and is required for the stability of newly translated Cox2p

**DOI:** 10.1101/2023.06.12.544600

**Authors:** Maria Antônia Kfouri Martins Soares, Letícia Veloso Ribeiro Franco, Jhulia Almeida Clarck Chagas, Fernando Gomes, Mário H. Barros

**Author notes:** Correspondence address: Mario H. Barros, Departamento de Microbiologia, Instituto de Ciências Biomédicas, Universidade de São Paulo, Av. Professor Lineu Prestes, 1374, 05508-900, São Paulo, Brazil;.

## Abstract

Based on available platforms of *Saccharomyces cerevisiae* mitochondrial proteome and other high throughput studies, we identified the yeast gene *DMO2* with a profile of genetic and physical interactions that indicate a putative role in mitochondrial respiration. Dmo2p is a homolog to human DMAC1 with two conserved cysteines in a Cx_2_C motif. Here, we localized Dmo2p in the mitochondrial inner membrane with the conserved cysteines facing the intermembrane space. The observed phenotypes of the *dmo2* null mutant indicate a general function in cellular stress response; the mutant displayed poor growth on non-fermentative media at 37°C, and in oleate, it is more sensitive to heat and oxidative stress while its overexpression confers resistance to some of the tested stressors. Dmo2p topology and modeled structure suggested a functional redox role for the Cx_2_C motif sustained by site-directed mutagenesis of both cysteine residues. The respiratory deficiency of *dmo2* mutants at 37°C led to a reduction in cytochrome *c* oxidase activity (COX) and the formation of *bc*1-COX supercomplexes; we also observed a rapid turnover of Cox2p, the subunit two of the cytochrome *c* oxidase complex that harbors the binuclear Cu_A_ center. Moreover, Dmo2p co-immunoprecipitated with Cox2p and components of Cu_A_ center maturation such as Sco1p and Sco2p; and finally, *DMO2* overexpression can suppress *cox23* respiratory deficiency, a mutant that has the mitochondrial copper homeostasis impaired. Overall, our data suggest that Dmo2p is required for Cox2p maturation, potentially by aiding proteins involved in copper transport and incorporation into Cox2p.

## Introduction

The mitochondrial intermembrane space (IMS) is a tiny sub-compartment localized between the outer and the inner membrane of mitochondria that plays crucial roles in many cellular processes, including the regulation of transport of proteins, lipids, and metal ions such as copper. The IMS also serves as a platform for redox signaling, whereby the redox state of proteins and metabolites within the IMS can influence cellular signaling pathways, energy production, and apoptosis. (Herrmann and Riemer, 2010).

Here, we detail the involvement of the poorly characterized protein Dmo2p in redox-dependent processes in the mitochondrial intermembrane space. *DMO2* has a homolog in humans, DMAC1, found in quantitative proteomic studies as required for the distal portion of the respiratory complex I membrane arm and supercomplex formation (Stroud et al., 2016).

According to the modeled structure and our localization data, Dmo2p is an integral inner membrane protein, displaying two conserved cysteines exposed into the intermembrane space (IMS). Indeed, most IMS proteins contain a conserved pattern of cysteine residues present in the “twin CX_3_C or CX_9_C motifs” (Vögtle et al., 2012) with a direct impact on the regulation of redox-sensitive cellular processes between the mitochondria and other compartments of the cell (Nuebel et al., 2016). Similarly to Dmo2p, many IMS products present dual localization (Yifrach et al., 2022); in this study, we corroborate these previous observations about Dmo2p localization.

Soluble and inner membrane-associated proteins in the IMS distribute copper to insert it into the Cox2p binuclear Cu_A_ center, Cox1p Cu_B_ site of COX. Additionally, they distribute copper to Sod1p the copper-zinc superoxide dismutase present in the cytosol and in the mitochondrial intermembrane space (Sturtz et al., 2001). Studies on *COX2* expression, its gene product translation, processing, and Cu_A_ formation unveiled several factors directly involved in this elaborate process (Fox, 2012; Jett and Leary, 2017). This intricate operation includes Cox17p, a soluble copper binding protein present in the cytosol and the IMS (Glerum et al., 1996a); Cox17p delivers copper to Sco1p (Horng et al., 2004), an integral inner membrane protein with its catalytic site facing the IMS where a Cx_3_C motif and a conserved histidine residue coordinate the metal ion; different residues of Sco1p have been found to specifically affect either Cox2 Cu_A_ site formation (Rigby et al., 2008) or Cox17-dependent copper addition (Cobine et al., 2006). Sco2p, Coa6p, and Cox16p are also described as participants of Cox2p-Cu_A_ site formation (Leary et al., 2009; Ghosh et al., 2016; Aich et al., 2018). Sco2p is not required for COX assembly itself but regulates copper efflux (Glerum et al., 1996b; Leary et al., 2007); Coa6p is also needed for the efficient formation of respiratory *bc*1-COX supercomplexes (Ghosh et al., 2016), while Cox16p has been detected in both Cox1p and Sco2p-Cox2p assembly complexes and has been proposed to be the assembly factor involved in the association of Cox1p and Cox2p assembly modules (Aich et al., 2018; Cerqua et al., 2018; Su and Tzagoloff, 2017). Finally, poorly characterized Cox23p, Cmc1p, Cmc2p, Coa4p, and Pet191p are soluble IMS proteins probably involved in copper delivery to COX and the regulation of copper trafficking (Barros et al., 2004; Horn et al., 2008; Horn et al., 2010; Swaminathan et al., 2022, Khalimonchuk et al., 2008). Here, we present evidence that Dmo2p also participates in the intricate process of Cox2p copper insertion.

## Results

### 1 Selection of DMO2 as a candidate to be studied

We have overexpressed uncharacterized genes that would unbalance the translation of mitochondrial products. First, we have cloned 32 putative components of the MIOREX complex (Kehrein et al., 2015) under the control of the strong *GAL10* promoter and evaluated the mitochondrial translation properties of the transformed yeast strain grown on galactose (Chagas et al., 2022); the overexpression of some of those genes causing a significant loss of the mitochondrial DNA includes the ORF *YDL157c*. Furthermore, we took advantage of the available information from the y3kproject.org (Stefely et al., 2016) and thecellmap.org (Usaj et al., 2017) to gather omic profiles of the *ydl157c* null mutant and the *YDL157c* gene product. It appeared to be a promising target for further study in both databases; in the y3kprojectwebsite, the *ydl157c* mutant showed a significantly lower abundance of proteins required for the OXPHOS system including Cox13p, Cox2p, Qcr8p, and Atp20p (y3k. project.org screenshot Figure S1A). Additionally, respiratory mutants such as *mrp10, shy1*, and *atp12* displayed a reduced abundance of the Ydl157c protein (Figure S1B); the genetic profile of the *ydl157c* mutant exhibited correlations with mutants encoding mitochondrial proteins and demonstrated genetic negative interaction with several mitochondrial mutants (thecellmap.org screenshot Figure S2A and S2B). Later, a machine learning multi-omic integration study confirmed our data mine and renamed *YDL157c* to *DMO2* (Determines Mitochondrial prOteome), describing its involvement in mitochondrial translation (Dickinson et al., 2022).

### 2 Phenotypic characterization of dmo2 mutants in different genetic backgrounds

Deletion of *DMO2* in the *S. cerevisiae* BY4742 strain background resulted in a respiratory growth defect phenotype at 37°C; on the contrary, in the W303-1A background, this phenotype was less pronounced (Figure 1A) at the indicated temperatures. These differences in respiratory phenotypes between the two backgrounds are due to a defective *hap1* mutant allele in BY4742. This allele affects the expression of several genes involved in mitochondrial biogenesis (Gaisne et al., 1999).

**Figure 1.**
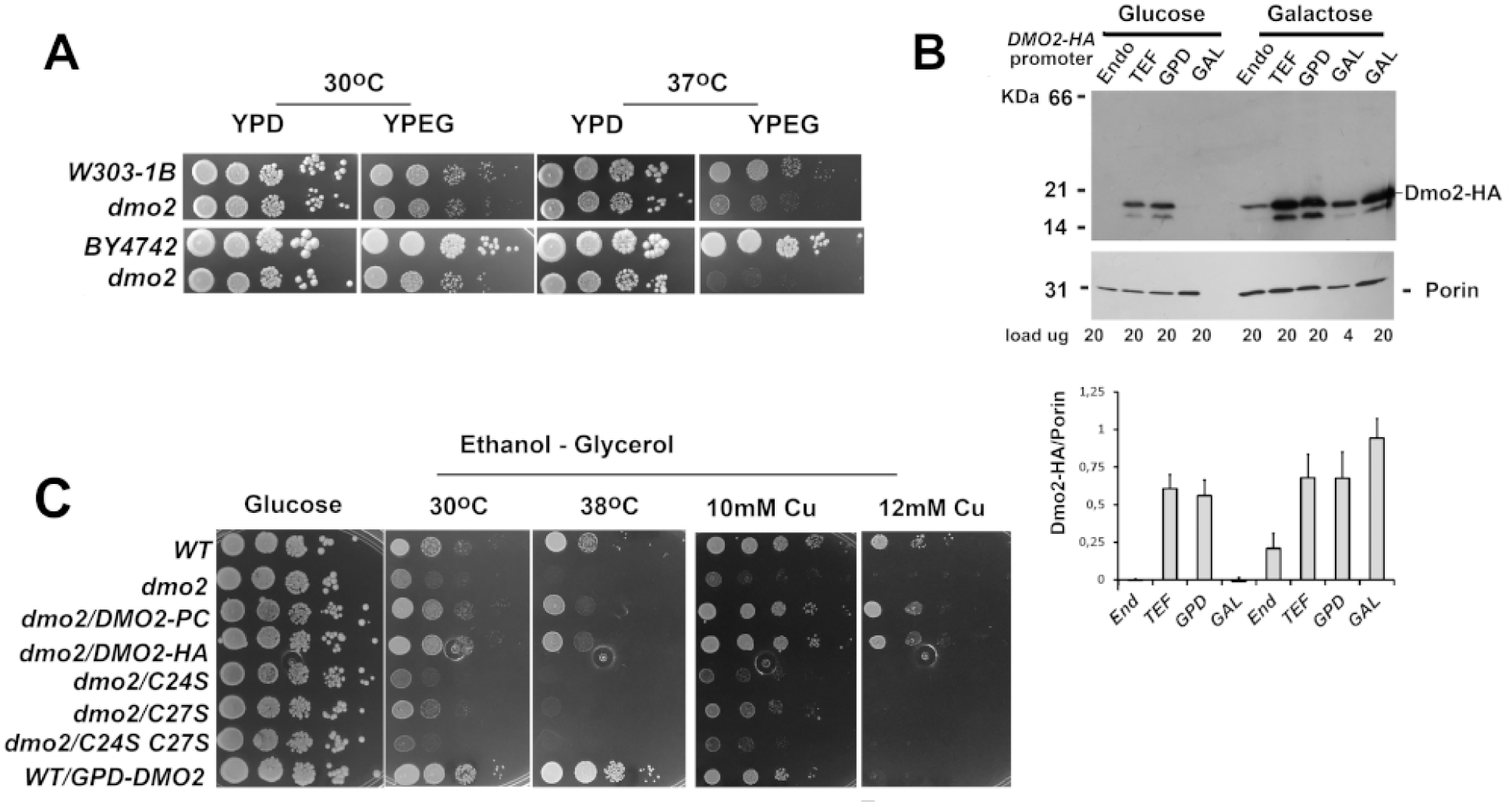
Growth properties of *dmo2* mutants. **(A)** Serial dilution growth test on rich fermentable glucose medium (YPD) and rich non-fermentable ethanol-glycerol medium (YPEG) incubated at 30°C and 37°C. Images related to W303-1B and BY4742 genetic background and the respective *dmo2* mutants (W303ΔDMO2-L, BY4742ΔDMO2). **(B)** Comparison of Dmo2p-HA protein levels after growth in glucose and galactose-rich media as indicated on the top. Mitochondria isolated from cells harboring *DMO2-HA* with the endogenous promoter (ENDO) and the *DMO2-HA* coding sequence under the control of TEF, GPD, and GAL promoters and their proteins separated in a 12.5% acrylamide gel by SDS-PAGE and immunoblotted against anti-HA and anti-porin antibodies as indicated. The bar graph below represents the average of two independent experiments with standard deviation in error bars. Band densitometry for Dmo2-HA/Porin ratio was calculated using the histogram component of GNU Image Manipulation Program GIMP 2.10.28. **(C)** Serial dilution tests on rich glucose medium (YPD) and rich ethanol-glycerol (EG) medium. The EG plates were also supplemented with 10mM CuSO_4_ and 12mM CuSO_4_ as indicated. Analyzed strains were BY4742 (wt) and BWΔDMO2 *dmo2* mutant transformed with the tagged alleles (*DMO2-HA*, *DMO2-PC*) the point mutants alleles *dmo2^C24S^*, *dmo2^C27S^*, *dmo2^C24S,C27S^*and the wild-type strain overexpressing *DMO2* under control of the *GPD* promoter (WT/GPD-DMO2).

We searched for distinct *dmo2* respiratory-deficient phenotype in the W303-1A background, with amenable possibilities to test its rescue by integrative plasmids at the *leu2*, *ura3,* and *trp1 loci*; therefore, the BY4742ΔDMO2 mutant was crossed to W303-1A and the resultant diploid sporulated; *dmo2* spores were selected and backcrossed to W303-1A five times successively to get the desired auxotrophies together with the typical W303 alleles; the final selected *dmo2* mutants transformed with tagged versions of the gene and the cysteine C24 and C27 mutants (Figure S3A); the protein C (PC) and hemagglutinin A (HA) tags inserted at the C-terminus of *DMO2* complemented the *dmo2* mutant phenotype (Figure 1C). The presence of a conserved cysteine pair present in the N-terminus (Figure S3B S3C) provided a hint linking Dmo2p to redox processes through thiol/disulfide exchange; thiol groups play crucial roles in the folding, stability, and activity of numerous proteins, including chaperones with distinct thiol switches mediating redox signaling (Demasi et al, 2021, Fra et al., 2017). To explore the significance of the cysteines to Dmo2p function, we replaced C24 and C27 for serines, leading to the single mutants *dmo2^C24S^*, *dmo2^C27S^*, and also the double mutant *dmo2^C24SC27S^*; growth properties of the new strains were evaluated on non-fermentable carbon sources media at 30°C and 38°C, and in the presence of 10mM or 12mM CuSO_4_. While the tagged *DMO2-HA* and *DMO2-PC* fusions completely restored the respiratory growth properties of the wild-type control, the cysteine mutants did not. Additionally, the overexpression strain with the *GPD-DMO2* construct grew better than the wild-type at 38°C (Figure 1C). Interestingly, the C27S mutant growth is slightly better than the null mutant and the one harboring the C24S mutation, particularly in the 10mM copper-containing medium. In the presence of 12mM CuSO_4_, the growth of the strain overexpressing *DMO2* was also severely affected. We then compared the survival of the wild-type strain, the *dmo2* mutant, and the strain overexpressing *DMO2* after a 5-hour challenge with different metal salts and oxidants (Figure 2A, Figure S3C). In this test, the *DMO2* overexpression resulted in lower survival capacity when challenged with copper, in agreement with its impairment growth in the copper media (Figure 1C). Interestingly, compared to the wild type and the overexpression strain, the null mutant displayed an increased formation of colonies resistant to the challenge with ZnCl_2_ (Figure S4A).

**Figure 2.**
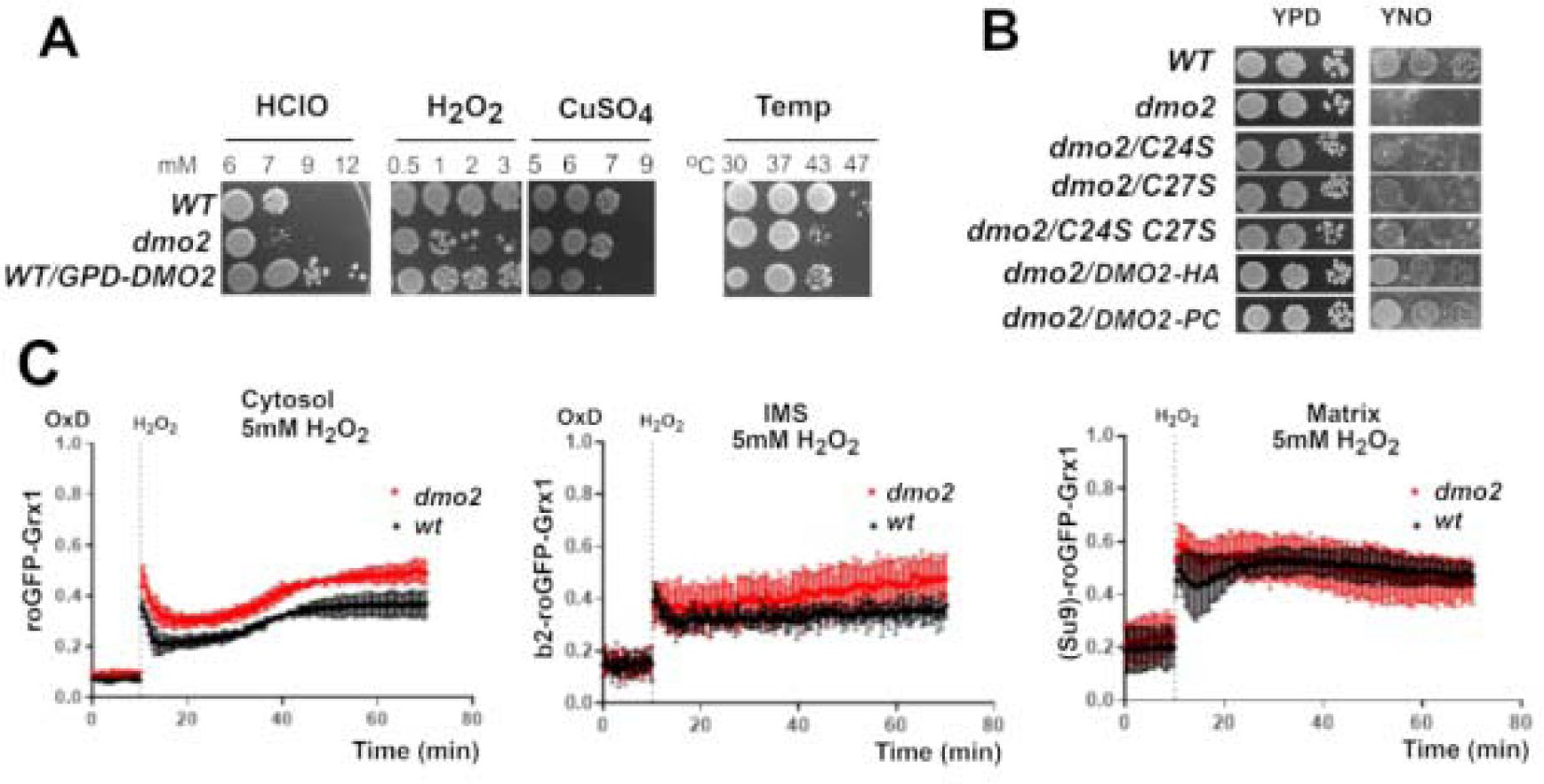
Stress tolerance response of the *dmo2* mutant. **(A)** Test of hypochlorite (HClO), hydrogen peroxide (H_2_O_2_), CuSO_4_, and temperature challenges. Equal amounts of cells from the wild-type strain W303-1A (wt), *dmo2* mutant (W303ΔDMO2-L), and the strain overexpressing *DMO2* under *GPD* promoter (W303+pCGPD-DMO2) were incubated for 5 hours in different concentrations of hypochlorite, hydrogen peroxide, CuSO_4_ and for 2 hours at the indicated temperatures (right panel). After the cells challenge, they were spotted on rich glucose media and grown at 30°C for two days. The complete panel for HClO, H_2_O_2_ and CuSO_4_ challenge is shown in Figure S4A. **(B)** Serial dilution test in glucose (YPD) and oleate (YNO) media of the W303-1A (wt), *dmo2* null mutant (*dmo2*), *dmo2* harboring a C24S point mutation (*dmo2/C24S*), *dmo2* with a C27S point mutation (*dmo2*/*C27S*), *dmo2* with both point mutations (*dmo2*Δ*/C24S C27S*) and *dmo2* harboring tagged versions of *DMO2* containing the hemagglutinin A or protein C epitopes (*dmo2/DMO2-HA, dmo2/DMO2-PC*). Plates were incubated at 30°C for three days and photographed. **(C)** Measurement of the cytosolic, intermembrane space (IMS), and mitochondrial matrix glutathione oxidation state was performed as described in the methods section. Values shown in red correspond to the *dmo2* null mutant, while values shown in black correspond to the parental W303-1B strain.

With the *DMO2-HA* construct, we evaluate the level of *DMO2* expression under the control of different promoters; the *GAL10-DMO2-HA* fusion showed higher levels of Dmo2p-HA compared to other constructs when cells were grown on galactose (Figure 1B), but it led to about 50% of petite colonies formation after overnight culture, with the frequency of petites estimated by crosses with ρ^0^ tester strains; the expression reached by the *GPD-DMO2* construct did not compromise the mtDNA stability; moreover, the toxicity of the *GAL-DMO2* construct led to a retard in galactose growth (Figure S4B).

To gather additional phenotypic information on *DMO2*, we challenged the null mutant and an overexpression strain with different stressful conditions. The *dmo2* null mutant was more susceptible to hypochlorite, hydrogen peroxide, and temperature stresses (Figure 2A, Figure S4A); the higher sensitivity to high temperature led us to investigate whether cells lacking *DMO2* could adapt to heat shock responses, as demonstrated in Susan Lindquist’s classic experiment (Yost and Lindquist, 1991). Wild-type cells and *dmo2* mutants were incubated at 37°C for 15 minutes, while another set of the same mutants was kept at 23°C. As expected, wild-type cells exhibited higher resistance to subsequent heat shock treatments, whereas the *dmo2*Δ mutant did not (Figure S5), indicating a potential thermal stress protection conferred by Dmo2p. When spotted on a non-fermentable oleate medium, the *dmo2 null* mutant did not grow, the point mutants *dmo2^C24S^*, *dmo2^C27S^*, *dmo2^C24S,C27S^* showed poor growth, and the tagged genes *DMO2-HA* and *DMO2-PC* rescued the growth capacity of the wild-type strain (Figure 2B).

While the *dmo2* null mutant was more sensitive to hypochlorite, its overexpression reached by the *GPD-DMO2* fusion (Figure 1C) provided protection when challenged with hypochlorite. However, it exhibited elevated sensitivity to copper (Figure 2A, Figure S3C). Hypochlorite is a powerful oxidant that induces oxidation of numerous biological molecules including the α-amino groups of His, Arg, Lys, the sulfured Cys and Met, and the aromatic Trp and Tyr (Hawkins and Davies, 2019); it also causes dose-dependent depletion of reduced glutathione and apoptosis induction in yeast (Kwolek-Mirek et al., 2011). Therefore, the increased sensitivity to hypochlorite prompted us to evaluate the redox state of the *dmo2* mutant strain by measuring the redox state of the cytosolic and intramitochondrial glutathione pools (Figure 2B). Glutathione plays a crucial role in yeast by reducing disulfide bonds and protecting cysteines from over-oxidation (Grant et al., 1996). By using a genetically encoded fluorescent probe sensitive to the GSH: GSSG redox couple (Grx1-roGFP2) (Hanson et al., 2004), we observed a slightly higher content of oxidized glutathione (GSSG) in the cytosol and mitochondrial intermembrane space of the *dmo2* null mutant compared to wild-type cells under the same oxidant conditions, while in the mitochondrial matrix, there was not a significant difference in both strains (Figure 2C).

Altogether, the *dmo2* mutants have growth deficiency on non-fermentable carbon sources at 37°C, accompanied by raised sensitivity to heat stress. It also presents poor growth on oleate and increased sensitivity to hypochlorite and hydrogen peroxide stress, followed by higher levels of oxidized glutathione in the cytosol and in the intermembrane space; the overexpression of *DMO2* was mainly protective to hypochlorite but sensitive to copper. Therefore, the *dmo2* phenotypic characterization sustains a role for Dmo2p in cellular stress response, possibly in a redox-dependent manner.

### 3 Dmo2p intracellular localization and distribution

Proteome-profile studies detected Dmo2p on highly purified mitochondria (Reinders et al., 2006). The tagged Dmo2-HA enables us to follow its submitochondrial distribution and solubilization properties after sonication of mitochondria with or without sodium carbonate. Additionally, it allows us to assess its distribution in mitochondria and mitoplasts subjected to proteinase K digestion. Sodium carbonate did not solubilize Dmo2p, confirming its predicted membrane association as indicated by its alpha-fold modeled structure (Jumper et al., 2021), which suggests the presence of two transmembrane domains (Figure 3A and Figure S3A); moreover, the protein is susceptible to proteinase K after the osmotic removal of the outer membrane, indicating that its C-terminus is an intermembrane space-exposed domain (Figure 3B). The outer-membrane removal by hypotonic shock was confirmed by the substantial loss of the soluble intermembrane marker cytochrome *b2* and supported by the decrease of Sco1p protein levels as a result of the proteinase K digestion; therefore, Dmo2p has the same topology as the one described for Sco1p (Glerum et al. 1996b). The α-ketoglutarate dehydrogenase was protected against proteinase K in all conditions, confirming the intactness of the inner membrane in the mitoplasts (Figure 3B). We conducted additional subcellular fractionation experiments to evaluate the localization of Dmo2p in oleate-grown cells. Oleate is an oxidative medium known to induce peroxisome biogenesis and activity (Venhuis et al., 1987). This experiment was motivated by previous findings that suggested the presence of Dmo2p associated with peroxisome membranes or membrane periphery (Yifrach et al., 2022). In yeast, peroxisomes carry out the β-oxidation of fatty acids to generate acetyl-CoA, which produces high levels of reactive oxygen species (ROS) as byproducts; indeed, mitochondria and peroxisome play central roles in the metabolism of ROS, with a common evolutionary origin (Bolte et al., 2015), they also share components of the fusion-fission machinery (Fransen et al., 2017). Differential centrifugation of oleate-grown cells led to an enriched organelle pellet, which was subjected to ultra-centrifugation in a discontinuous sucrose gradient to separate mitochondria (top fractions) from peroxisome (bottom fractions) (McCammon et al., 1990). Mitochondrial Porin and Dmo2p-HA with a size below 21.5 KDa were distributed in the top fractions, while Dmo2-HA with a size higher than 31 kDa and Fox3p were mainly concentrated in the bottom (Figure 3C). Fox3p is a protein with a dual localization in mitochondria and peroxisome, so its distribution in the top fractions is also expected (Erdmann, 1994; Vögtle et al., 2012). Fox3p cleaves 3-ketoacyl-CoA into acyl-CoA and acetyl-CoA during the β-oxidation of fatty acids (Mathieu et al., 1993).

**Figure 3.**
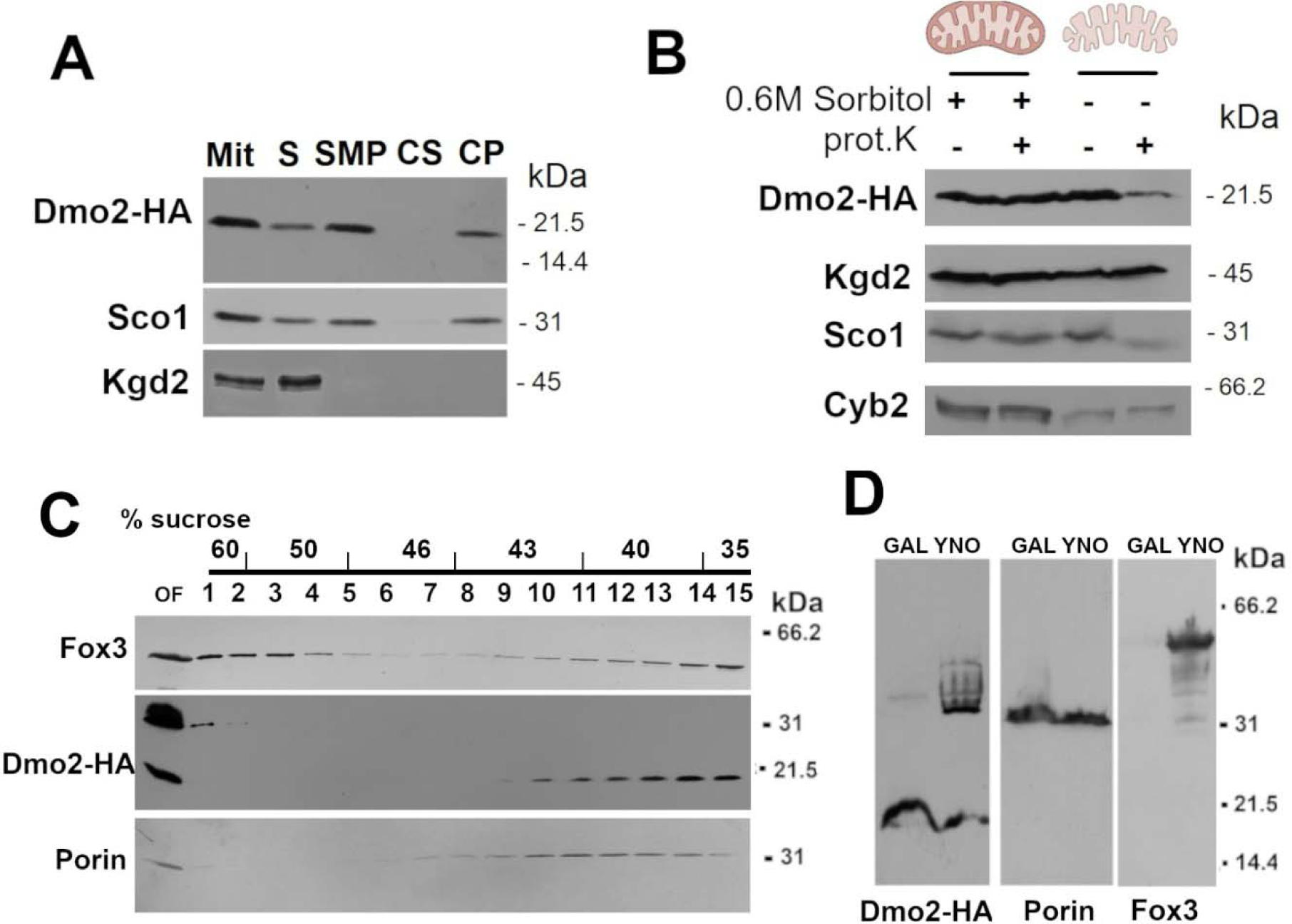
Dmo2p is located in the mitochondrial inner membrane facing the intermembrane space. **(A)** Membrane solubility assay. Mitochondria (Mit) isolated from the W303/*DMO2*-HA strain were sonicated for 10 seconds at 80% amplitude and centrifuged at 30,000x*g* for 30 minutes. The supernatant (S) and part of the pellet (SMP) were collected; the pellet was suspended in 0.2M sodium carbonate, incubated on ice for 30 minutes, and centrifuged again. The supernatant (CS) and pellet (CP) were collected. All fractions (Mit, S, SMP, CS, and CP) were separated in a 12.5% acrylamide gel by SDS-PAGE, transferred to a nitrocellulose membrane, and immunoblotted with anti-HA, anti-Sco1, and anti-Kgd2. **(B)** Intramitochondrial localization of Dmo2. Mitochondria from the W303/*DMO2*-HA strain were osmotically challenged with HEPES without sorbitol, generating mitoplasts. Mitochondria and mitoplasts were submitted or not to treatment with Proteinase K for 1 hour to test whether proteins are available for Proteinase K digestion. **(C)** The organelle fraction (OF) isolated from oleate-grown cells was subjected to ultracentrifugation in a discontinuous sucrose gradient as indicated in the panel; fractions were collected from top (15) to bottom (1) and subjected to western blot analyses with the antibodies indicated in the panel. **(D)** Proteins from organelle pellets obtained after growth of DMO2-HA cells in galactose (Gal) and oleate (YNO) were separated in a standard 12.5% acrylamide gel by SDS-PAGE and tested against the antibodies indicated at the bottom of each panel.

Dmo2-HA levels were also compared from organelle fractions (mitochondria and peroxisome) obtained from cells grown on galactose and oleate; in the oleate protein extracts, the Dmo2-HA with lower mobility was abundant, while in the galactose extract, only a faint band can be distinct in the same region (Figure 3D). Fox3p, similarly, was most detected in the extracts obtained from oleate-grown cells. Porin and Dmo2-HA, with the expected size, were equally detected in both extracts.

To test whether the unexpected high molecular weight bands observed for Dmo2p were due to disulfide bond formation in the oxidative oleate medium, we examined if the cysteine point mutants C24S and C27S variants also display the same mobility behavior of Dmo2-HA in high molecular weight bands; indeed, in organelle pellets from cells grown on oleate medium, all Dmo2-HA protein versions displayed a high molecular weight product; but in galactose the Dmo2-HA products migrated with the expected mobility (Figure 4a); in conclusion, the higher molecular weight bands in the double mutant C24S-C27S excluded the formation of a disulfide-bonded intermediate of Dmo2p in this mutant. Interestingly, the stability of the different Dmo2-HA products varied across the tested media; while the mutants harboring the C24S were stable in galactose-grown cells, they seemed less stable in the oleate medium, and only the product with higher molecular weight was detected; the detection of the C27S product was similar to that observed for the wild-type Dmo2p-HA allele except that the protein seemed less stable in the extract obtained from galactose-grown cells.

**Figure 4.**
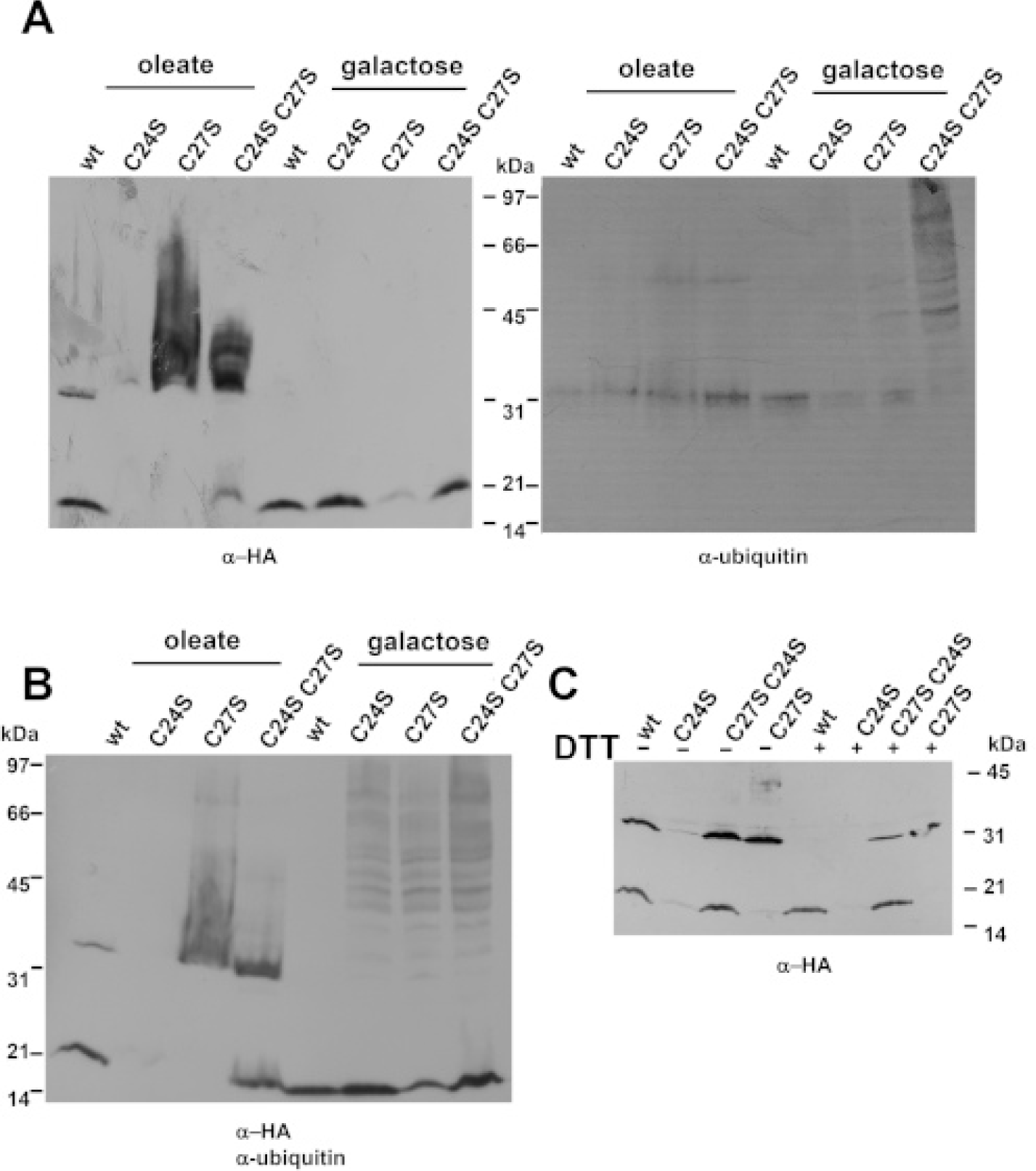
Dmo2p protein size variation. **(A)** The organelle fraction – containing both mitochondria and peroxisomes) extracted from cells containing different *DMO2* alleles (the wild-type *DMO2* (wt), and the point mutants C24S, C27S, and C24S-C27S double mutant) were grown on galactose and oleate. The protein extraction was immunoblotted against anti-HA, anti-ubiquitin, or both **(B)**. **(C)** The organelle fraction obtained from oleate growth from the same strains was treated (+) or not (-) with 100mM DTT and heat for 5 minutes at 100°C and immunoblotted against anti-HA.

We also examined whether Dmo2-HA high molecular weight bands were susceptible to treatment with a very high concentration of DTT (100mM) and heat. In the Dmo2-HA strain, the aberrant products disappeared after the DTT treatment, the same was true for C27S, and for the double mutant C24S C27S - although less intense - the aberrant bands were still detected after DTT addition; curiously, the very high concentrations of DTT and heat led to the disappearance of the abnormal band in the wild-type, but the expected Dmo2p product was not increased after this treatment, indicating that it was completely degraded; we then reasoned that the higher molecular weight bands observed for Dmo2p may have originated from post-translational modifications or Dmo2p aggregates induced under oxidative stress; at any rate, the introduction of the C27S mutation leads to a higher accumulation of the altered versions of Dmo2p; we inferred that a plausible modification that fits the pattern of the ladder bands observed in this situation is ubiquitination; mitochondrial proteins are ubiquitinated in yeast when exposed to the cytosolic surface of the outer membrane (Lehmann et al., 2016), which makes sense for Dmo2p, considering its proposed to transit towards the peroxisome under redox stress, getting, in this traffic, the ubiquitin modifications (Sugiura et al., 2017). Therefore, we tried out an anti-ubiquitin antibody (P4D1) on the same protein extracts. Although we could not observe a perfect coincidence in band distribution that would explain the different molecular weight bands in Dmo2-HA, it is noticeable that in the strain expressing the wild-type *DMO2-HA,* there is less reaction of the anti-ubiquitin antibody in comparison to the *dmo2* cysteine mutants, indicating a rise in ubiquitinated proteins in the absence of wild-type *DMO2* in galactose-grown cells.

In conclusion, Dmo2p excess resulted in increased resistance to hypochlorite (Figure 1C) while *dmo2* mutants showed increased sensitivity; *dmo2* mutants grow poorly in oleate, and in this medium, Dmo2p also displayed a high molecular weight product that is enriched in the peroxisome fraction. Overall, our data supports the hypothesis that the two cysteines function on thiol-based mechanisms. They are needed for proper response and adaptation to altered redox states. However, we could not determine the nature of the aberrant Dmo2p products, especially those present in the C27S mutants. We speculate that the C27S mutation leads to conformational changes inducing aggregation.

### Dmo2 role in mitochondria respiration

Considering the observed growth defects of *dmo2* cells on non-fermentable carbon sources at 37°C, the mtDNA instability under *GAL-DMO2* overexpression, and its reported deficiency in mitochondrial translation (Dickinson et al., 2022), we evaluated the respiratory properties of *dmo2* mutants in the W303 background, starting with OXPHOS complexes activities, and oxygen consumption in mitochondria isolated from wild-type and *dmo2* cells grown on galactose media at 30°C and 37°C. In agreement with their growth properties on non-fermentable media (Figure 1), lower oxygen consumption and OXPHOS activity were observed in mitochondria isolated from the mutant *dmo2* at 37°C; cytochrome *c* oxidase activity (CIV) was particularly affected under these conditions (Figure 5). Mitochondria obtained from *dmo2* cells at 37°C also showed reduced steady-state levels of ATP synthase dimers in comparison to the wild-type mitochondria obtained from cells also grown at 37°C. Previous proteomic data from the *dmo2* mutant displayed a reduction in Atp20p, a protein required for ATP synthase dimerization (Reinders et al., 2007; Stefely et al., 2016) (Figure S1). Furthermore, the steady-state level of the 1[COX]2[bc1] supercomplexes were decreased in the *dmo2* mutants as observed using anti-Cox2 (Figure 5B), anti-Cor1, anti-Cob, and anti-Cox1 (Figure S7). Therefore, we concluded that the *dmo2* deletion impaired respiration at 37°C, with complex IV being more severely affected.

**Figure 5.**
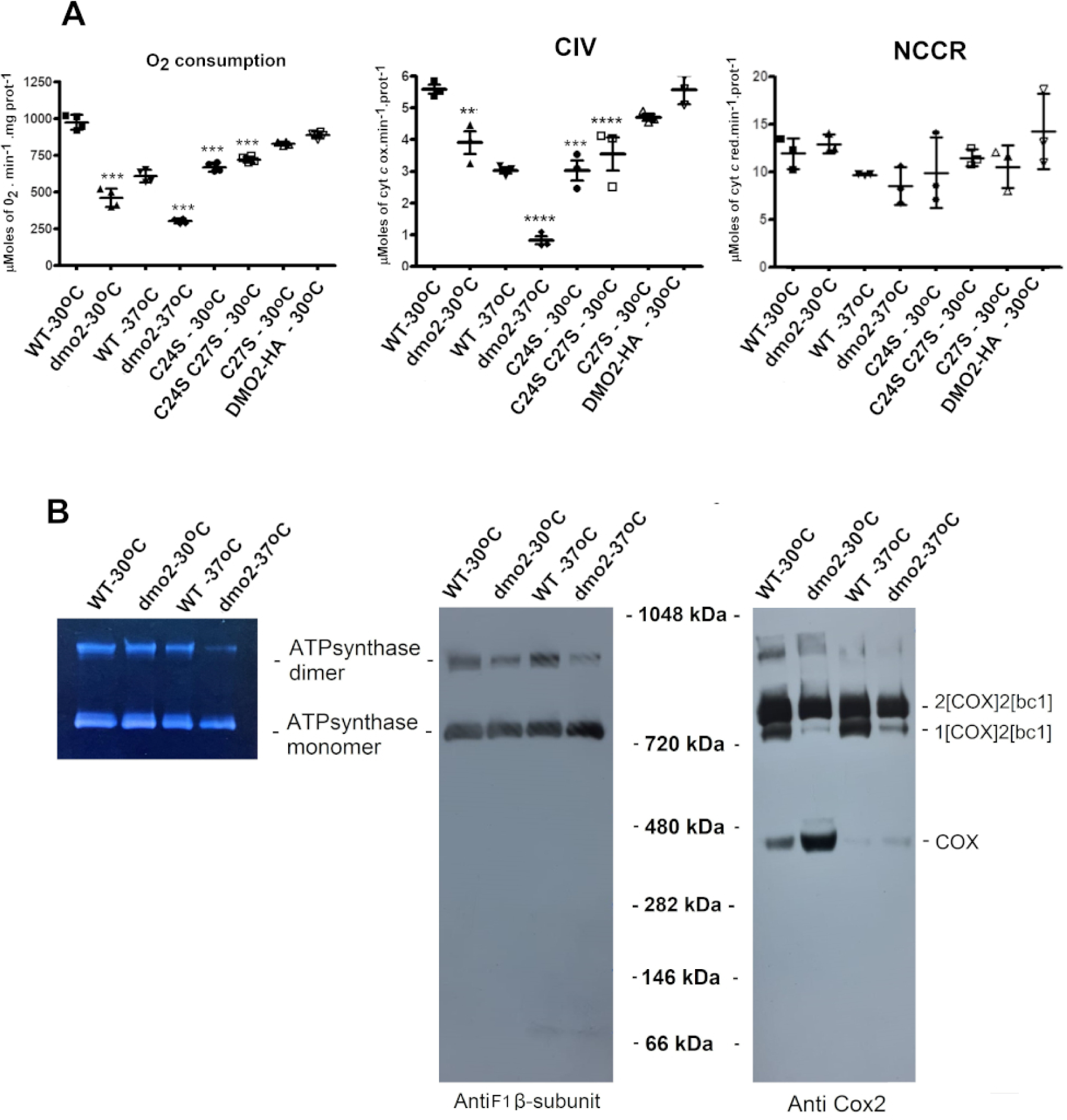
Respiratory properties of *dmo2* mutants. The indicated assays were performed with mitochondria isolated from W303-1A (wt) and W303ΔDMO2-L (*dmo2 null mutant)*), the *dmo2* point mutants C24S, C27S, C24SC27S and the *dmo2* mutant expressing the *DMO2-HA* construct; mitochondria obtained from cells grown on galactose at 30°C, and 37°C and assayed as indicated in the panels **(A)** Respiratory capacity was measured by oxygen consumption after NADH addition using a Clark electrode as shown in the panel on the left; the NADH dehydrogenase and CIII activities (NCCR) was assessed by the rate of cytochrome *c* reduction, while cytochrome c oxidase activity (CIV) by the rate of reduced cytochrome *c* oxidation. Data were obtained by three independent measures and analyzed in GraphPad Prism (La Jolla, CA, USA) ***p<0.001, ****p<0.01, **p<0.05 versus WT. **(B)** Isolated mitochondria were extracted with 2% digitonin and loaded into a non-denaturing 4-13% acrylamide gradient gel by BN-PAGE. The gel on the left illustrates ATPase in-gel activity assay as described in the methods section. For the middle and right panels, proteins separated in the gel were transferred to a PVDF membrane and probed with anti-subunit β of F_1_ ATP synthase (middle) and anti-Cox2 (right).

Mitochondria translation defects cause pleiotropic deficiencies in the OXPHOS complexes (Tzagoloff and Dieckmann, 1990). Based on the reported involvement of Dmo2p in mitochondrial translation (Dickson et al., 2022), we evaluated the turnover of newly synthesized mitochondrial products in *dmo2* cells grown in galactose. After a 20-minute *in vivo* labeling pulse at room temperature, we chased the radiolabeled turnover of the mitochondrial products for 20 and 40 minutes at 37°C. Under these experimental conditions, we did not observe a significant deficit in the accumulation of mitochondrial products, indicating that there was no general translation deficiency (Figure 6A). In our assays, the mutant *dmo2* exhibited higher turnover of the Cox2p product at both time points of chase (Figure 6B); indeed, when compared with wild-type mitochondrially encoded products, only Cox2p turnover was significantly increased. The elevated turnover of Cox2p in the mutant did not result in a significant change of its steady-state level in comparison with the wild type. This was also observed for other mitochondrial proteins (Figure S8A), nor did the addition of *yme1* deletion stabilize Cox2p in the *dmo2* background (Figure S8B). Yme1p is the catalytic subunit of the i-AAA protease complex responsible for the degradation of unfolded or misfolded mitochondrial gene products (Schreiner et al., 2012).

**Figure 6.**
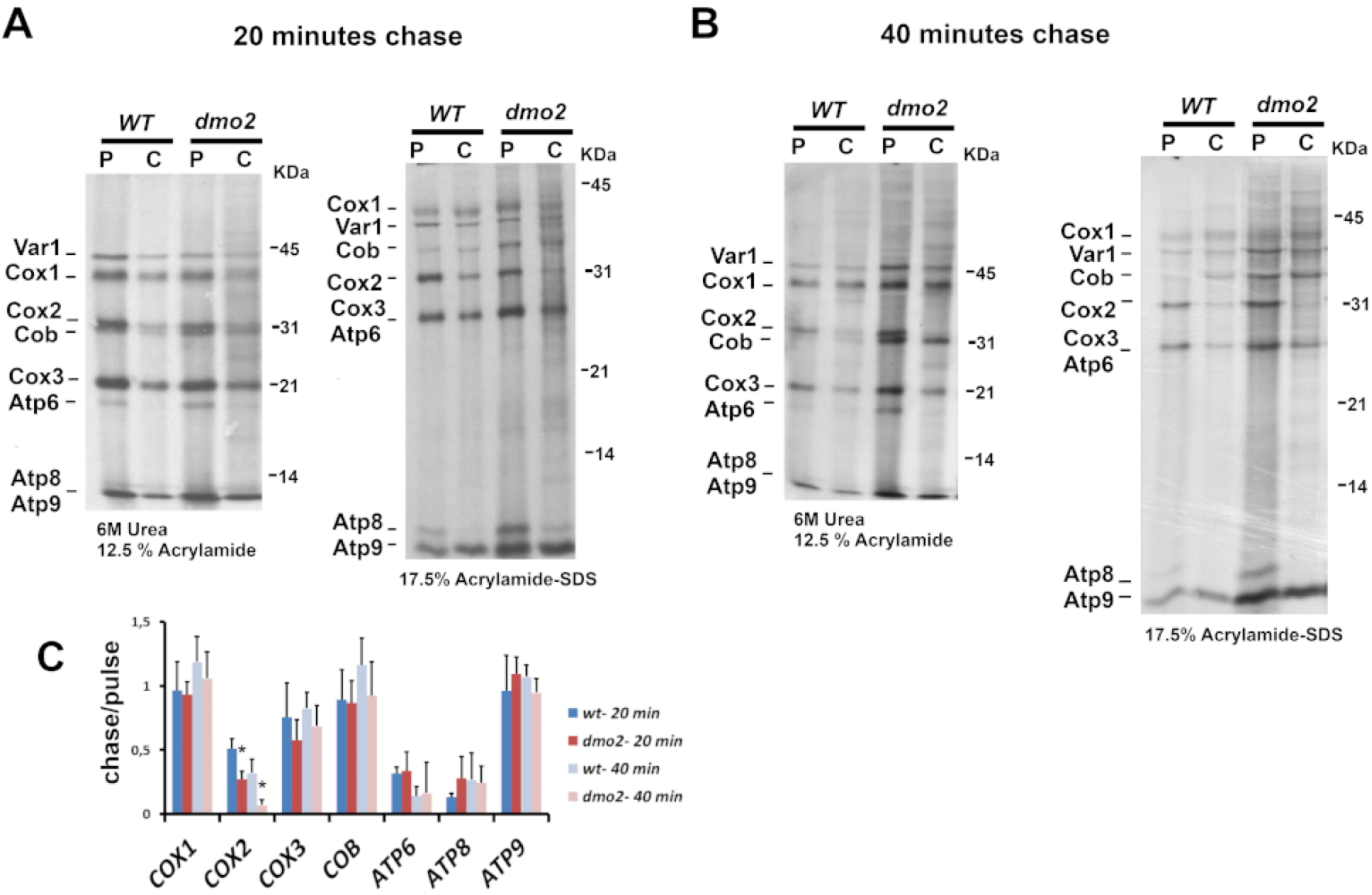
*dmo2* mitochondrial translation properties. *In vivo* labeling pulse and chase experiment. W303-1B (wt) and W303ΔDMO2-L (*dmo2*) were treated with cycloheximide for 10 minutes and incubated with ^35^S-methionine/cysteine mix for 20 minutes (pulse, P). Following that, methionine/cysteine mix was added, and cells were incubated for 20 minutes (Chase, C) (**A**) or 40 minutes (**B**). Radiolabeled mitochondrial products were separated and are indicated in the margins of the figure as follows: Cox1, Cox2, and Cox3 forming the catalytic core of cytochrome *c* oxidase; Atp6, Atp8, and Atp9 components of the ATP synthase; Cob, a constituent of the bc1 complex; mitoribosome protein Var1. The graphs below represent the average chase and pulse ratios of band intensity count measurements obtained from four independent labeling experiments. The bars are standard deviations of bands densitometry counts obtained using the histogram component of GNU Image Manipulation Program GIMP 2.10.28.* p< 0.05 vs WT.

### Genetic interactions of DMO2 with Cox2p processing factors

As mentioned above, the Cox2p maturation process includes the addition of two copper ions at the Cu_A_ site of the protein that can be associated with *dmo2* phenotypes; we hypothesized that if Dmo2p functions in the Cox2p metal chaperone complex, it may exhibit physical or genetic interactions with genes implicated in Cox2p module assembly (Franco et al., 2018).

Translation of *COX2* mRNA depends on Pet111p activation (Mulero and Fox, 1993); the resulting Cox2p is co-translationally inserted in the inner mitochondrial membrane (IMM), a process mediated by Oxa1p and Mba1p. Cox20p is a chaperone that stabilizes and facilitates membrane insertion of newly synthesized Cox2p, protecting it from degradation (Hell et al., 2000; Eliot et al., 2012; Bourens et al., 2014; Lorenzi et al., 2016; Bourens and Barrientos, 2017); after insertion of the N-terminal domain in the IMM, a complex comprising Imp1p, Imp2p, and Som1p cleaves the hydrophilic N-terminal portion of Cox2p (Nunari et al., 1993; Jan et al., 2000). Subsequently, the second transmembrane domain, the C-terminal portion of Cox2p, is then inserted in the IMM with the aid of Pnt1p, Mss2p, and Cox18p (Saracco and Fox, 2002). As a result of this process, Cox2p is inserted in the IMM, chaperoned by Cox20p, and ready to receive two copper ions in the Cu_A_ site (Fiumera et al., 2007; He and Fox, 1999; Broadley et al., 2001; Saracco and Fox, 2002). Sco1p and Sco2p, in a complex with Coa6p and Cox12p, deliver two copper ions to the Cox2p Cu_A_ site concomitantly to Cox18p dissociation (Bourens and Barrientos, 2017; Glerum et al., 1996a; Banci et al., 2008; Ghosh et al., 2016). Although still elusive, Cmc1p may play a role in copper delivery to Ccs1p, which then delivers it to Sod1p – and to Cox17p (Horn et al., 2008). Cox17p is a soluble intermembrane space (IMS) protein that delivers copper ions to both Sco1p and Sco2p (Glerum et al., 1996a, 1996b, Horng et al., 2004), while Cox23p – a homolog of Cox17p – has a putative function in copper trafficking as well (Barros et al., 2004); the process illustrated in Figure 7 depicts Cox2p translation, maturation, and copper addition processes.

**Figure 7.**
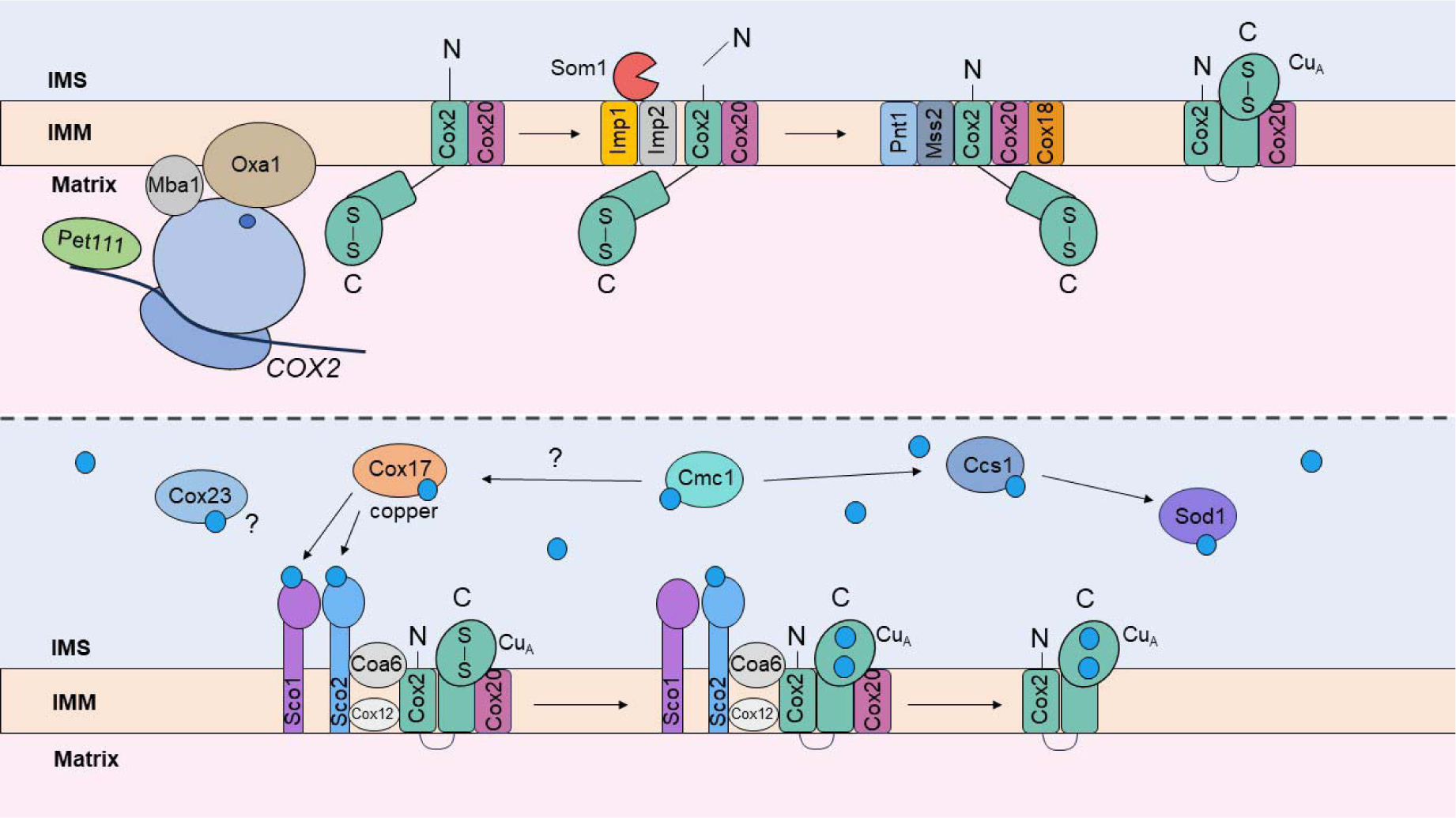
Cox2p translation, maturation, and copper addition process. The upper panel is relative to Cox2p insertion in the IMM; the lower panel is relative to copper insertion in the Cox2p Cu_A_ site. Adapted from Jett and Leary, 2017.

We searched for genetic synthetic deficiencies using the mutants in W303 genetic background for crosses. In the obtained double mutants, we noticed that *cox20 dmo2* retained the ability to revert to respiratory competency typically displayed in the *cox20* single mutant after a longer incubation time (Figure S6); however, the double mutant *cox20 dmo2* seems to revert at a lower frequency in comparison to the single *cox20* mutant (Figure S6). The *cox20* mutants can recover their respiratory deficiency through a bypass promoted by *yme1*, *mgr1*, or *mgr3* mutations (Elliot et al., 2012).

The *sco2* mutant grows as well as the wild-type cells in non-fermentable carbon sources (Glerum et al., 1996a), but the double *sco2 dmo2* showed diminished growth capacity on non-fermentable media (Figure 8A). This negative interaction between the two mutants is consistent with high throughput synthetic genetic array studies (Figure S2; Usaj et al., 2017), and it makes sense with the proposed function of Dmo2p in copper homeostasis as Sco2p regulates mitochondrial copper efflux (Leary et al., 2007).

**Figure 8.**
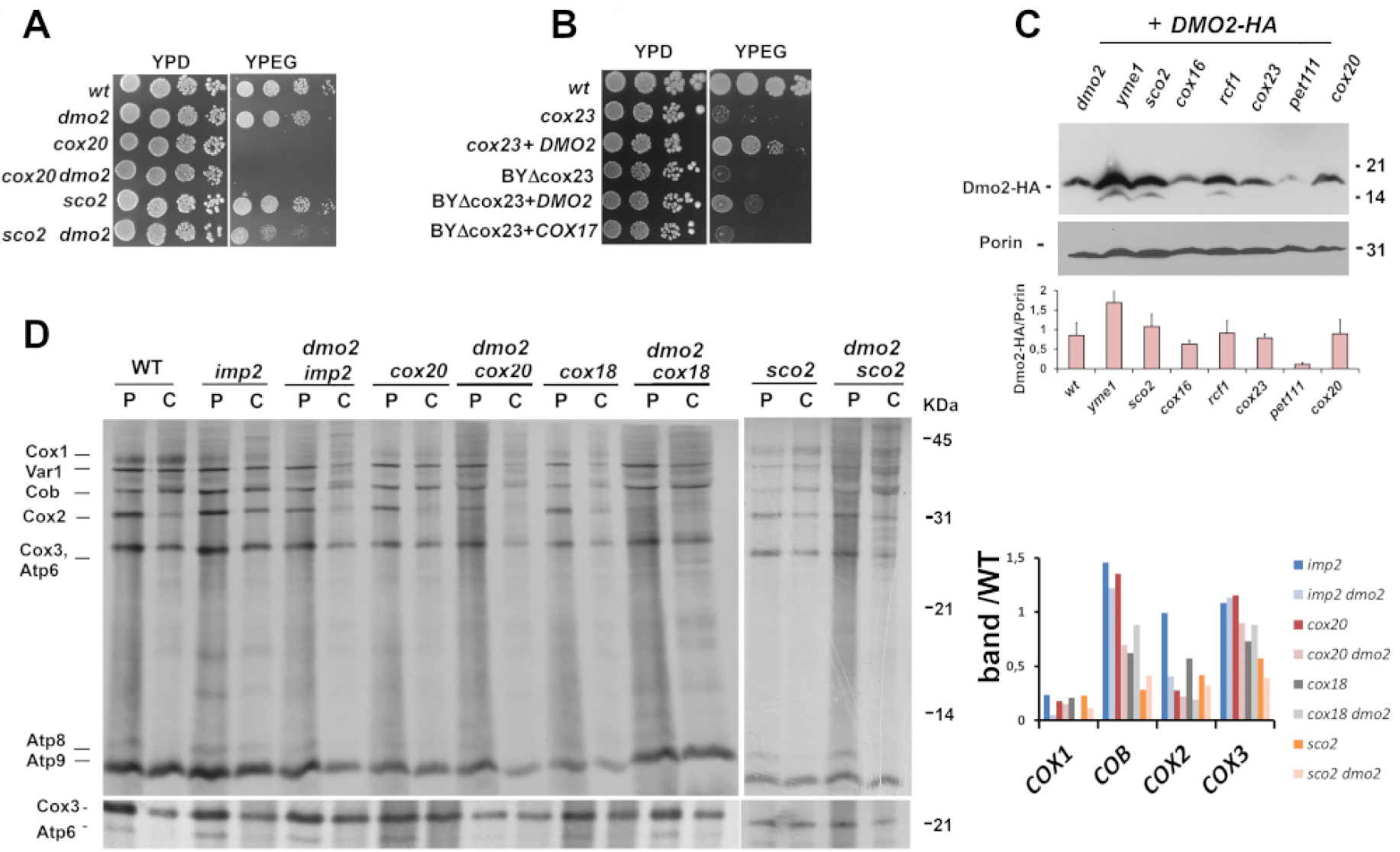
Overexpression of *DMO2* partially rescues *cox23* growth. **(A)** Growth of yeast strains after serial dilutions in rich glucose media (YPD) and rich non-fermentable media (YPEG). The tested strains are the wild type (wt) W303-1A and derivative mutants: *dmo2*, *cox20*, *cox20 dmo2*, *sco2* and *sco2 dmo2*. Plates were incubated at 30°C or 37°C as indicated and photographed after two days of growth. **(B)** Serial dilution test in YPD and YPEG media of strains W303-1A (wt) and *cox23* harboring a YEp351-*DMO2* with *DMO2*’s endogenous promoter plasmid (*cox23* + *DMO2*). Growth of the *cox23* mutant in the BY4741 background (BYΔcox23) was also compared to the transformed mutant with *DMO2* (BYΔcox23+*DMO2*) and *COX17* (BYΔcox23+*COX17*). **(C)** Steady-state Dmo2p-HA protein levels in the indicated null mutants. The bar graph below represents the average of two independent immunoblots probed with anti-HA and anti-Porin, with error bars. **(D)** *In vivo* labeling pulse and chase. Cells were treated with cycloheximide for 10 minutes and incubated with ^35^S-methionine/cysteine mix for 20 minutes (pulse, P); followed by the addition of a mixture of cold methionine/cysteine with a new incubation for 45 minutes (Chase, C). Radiolabeled mitochondrial products were separated, as in Figure 6A, and are indicated in the margins of the figure. The band intensity ratio of the labeled Cox1p, Cox2p, Cox3p, and Cob products in each mutant by the wild-type correspondent product are plotted on the graph aside.

We further explored the genetic connection by transforming *dmo2* with various multi-copy plasmids harboring genes involved in Cox2p maturation, such as *SCO1, SCO2, COX17, COX18, COX19, COX23*, and the nuclearly encoded version of *COX2* (Supekova et al., 2010), which resulted in any growth improvement (data not shown). Conversely, we transformed the respective mutants for the same genes with *DMO2* in a multi-copy plasmid (Figure S9); the over-expression of *DMO2* led to the suppression of the *cox23* respiratory growth deficiency in the two assessed parental backgrounds. (Figure 8B).

In further exploration, Dmo2p-HA steady-state level was evaluated in different mutants involved in Cox2p module formation and in other respiratory mutants. We detected a decrease of Dmo2p-HA content in *pet111* null mutant mitochondria. In the *cox23* and *cox16* backgrounds, the lower amount of Dmo2p-HA was not substantial, while in the *yme1* mutant, Dmo2p levels are very high (Figure 8C), as also detected in the y3k project (Stefely et al., 2016). Pet111p is essential for Cox2p translation, and Cox16p is crucial for the association between Cox1p and Cox2p assembly modules (Green-Wilms et al., 2001; Su and Tzagoloff, 2017).

The synthesis and turnover of mitochondrial products assessed in the double mutants *dmo2 imp2, dmo2 cox18, dmo2 cox20*, and *dmo2 sco2* unveil a general instability of the radiolabeled products. (Figure 8E). Atp6p and Atp8p were poorly detected, hampering proper comparisons. To follow the turnover of the mitochondrial products, we repeated the chase experiment; however, the higher instability of the double mutant products hampered the correct band identification after the chasing time. In this experiment, Cox1p translation was lower in *imp2 dmo2*, *cox18 dmo2*, and *sco2 dmo2* cells; similarly, Cox2p accumulation was shortening in these mutants but to a minor extent, except for *cox18 dmo2* (Figure 8D). In the *cox18* mutant, only the N-terminus domain of Cox2p can cross the inner membrane, with the C-terminus remaining exposed on the matrix side, and the copper atoms are not added (Fiumera et al., 2007); the removal of Cox2p N-terminus hydrophilic pre-sequence requires Imp2p, but it has no interference in the C-terminus copper site (Nunari et al., 1993). As expected, in the *cox20* mutant, the Cox2p product presents elevated turnover. Finally, while the *sco2* mutant did not show any particular deficiency, the double mutant *sco2 dmo2* not only displayed decreased stability of Cox2p but also showed a general impairment in mitochondrial translation (Figure 8D).

### Physical interaction of Dmo2p with Cox2p associated factors

We performed two forms of analysis to investigate whether Dmo2p physically associates with putative protein partners. First, we used dodecyl-maltoside to solubilize mitochondrial membranes embedded proteins, followed by ultracentrifugation in sucrose gradients to evaluate and size putative complex formation. In the control strain (*dmo2/DMO2-HA*) the Dmo2-HA product co-fractionated with the 144 kDa lactate dehydrogenase marker and lighter than porin sedimentation, which in blue native gel gives a product of around 480 kDa. We then hypothesized that Dmo2p may be associated with ancillary proteins of Cox2p, such as Sco1p and Sco2p, which share with Dmo2p their localization and topology. Taking advantage of available antibodies (kindly provided by A. Tzagoloff – Columbia University), we observed co-sedimentation of Dmo2p with Sco1p and Sco2p (Figure 9); actually, Sco1p presents two peaks: the lighter one at fraction 12 may correspond to the native 88 kDa homotrimer form previously reported (Beers et al., 2002). We used anti-Prx1 (gift from M. Demasi – Instituto Butantan) as a sedimentation control of a mitochondrial antioxidant enzyme, which we have used before to follow lighter fractions in this type of sucrose gradient (Chagas et al., 2022). We then asked whether Dmo2p-HA would have its mobility altered in these gradients in different mutants’ backgrounds. Therefore, we transformed the double mutants *dmo2 sco2*, *dmo2 cox23*, *dmo2 pet111*, *dmo2 cox16*, *dmo2 pet191* with the *DMO2-HA* construct to follow the protein when its genetic interactors are missing (*cox23* and *sco2*), when Cox2p is not being translated (*pet111*), in mutants with involvement in Cox2p module maturation (*cox16*) and in *pet191*, whose wild type product also has the same topology of Dmo2p (Khalimonchuk et al., 2008); interestingly, the pattern of Dmo2p-HA sedimentation were similar in the tested strains, it consistently co-sedimented with Sco1p and Sco2p in all tested backgrounds (Figure 9), except in the *sco2*, in which Dmo2-HA is part of higher molecular weight complexes. We reasoned that Sco2p directly interact with Dmo2p, avoiding disulfide linkages within large oligomeric complexes. Moreover, Dmo2p-HA bands with migration above the 31kDa marker were noticed again in the mutants’ backgrounds with sedimentation in the heavier fractions of the gradients, showing that the modified Dmo2p-HA is present in large complexes.

**Figure 9.**
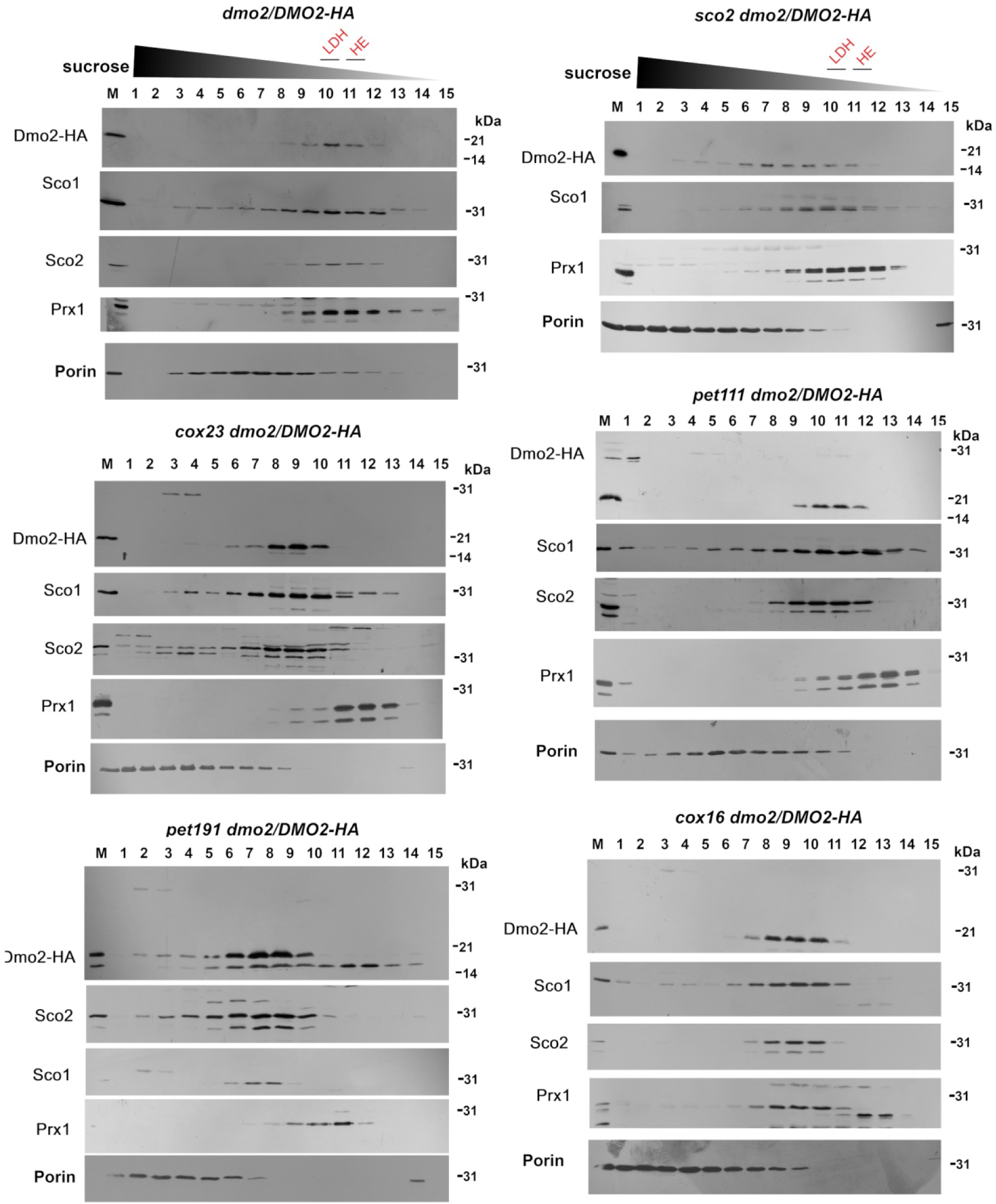
Dmo2 has a sedimentation pattern similar to Sco1 and to Sco2 in a sucrose gradient. Extracts from W303/DMO2-HA (*dmo2/DMO2-HA*), and the following mutants *sco2 dmo2/DMO2-HA*, *cox23 dmo2/DMO2-HA, pet111 dmo2/DMO2-HA, pet191 dmo2/DMO2-HA,* and *cox16 dmo2/DMO2-HA* were mixed with hemoglobin (64kDa) and lactate dehydrogenase (144kDa), applied on a 7-20% sucrose gradient, and ultra-centrifuged as described in the methods section. Gradient fractions were collected from bottom (1) to top (15) analyzed in a 12% acrylamide gel by SDS-PAGE. Proteins were transferred to a nitrocellulose membrane and probed with anti-HA, anti-Sco1, anti-Sco2, anti-Prx1, and anti-porin.

Another approach to investigate Dmo2p physical partners was conducted in co-immunoprecipitation assays using the construct *DMO2-CH* expressing Dmo2p with its C-terminus fused to a tandem tag of protein C epitope followed by polyhistidine; in these experiments, we used digitonin, a mild detergent, to extract mitochondrial proteins; Dmo2p-CH was immunopurified on protein C antibody beads, while the immunoblots of the eluate from the beads detected Dmo2p-CH, Sco1p, Sco2p, and Cox2p (Figure 10A), showing their co-immunopurification with Dmo2p-CH. In the eluate of the control strain harboring non-tagged Dmo2p, a faint band of Cox2p was detected, indicating that a small amount of Cox2p can interact non-specifically with the beads. In the non-tagged Dmo2p control eluate, Sco1p or Sco2p were absent. To confirm Dmo2p interaction with Cox2p, we transformed a yeast strain harboring tagged Cox2p-HAC with an integrative plasmid expressing DMO2-HA. The Cox2-HAC has a tandem tag of the hemagglutinin followed by protein C (Franco et al., 2018). In this experiment, the anti-HA antibody recognizes the Cox2p-HAC and Dmo2p-HA constructs, but the protein C antibody is specific to Cox2-HAC. In the eluate of Cox2p-HAC interactors, we observed a product displaying the same size as the Dmo2-HA construct, corroborating the interaction between the two proteins (Figure 10B); probably, the Dmo2p-HA in the eluate would increase if we were able to assess the co-immunoprecipitation promoted by the tagged *COX2-HAC* allele in the *dmo2* null mutant background transformed with the *DMO2-HA* fusion.

**Figure 10.**
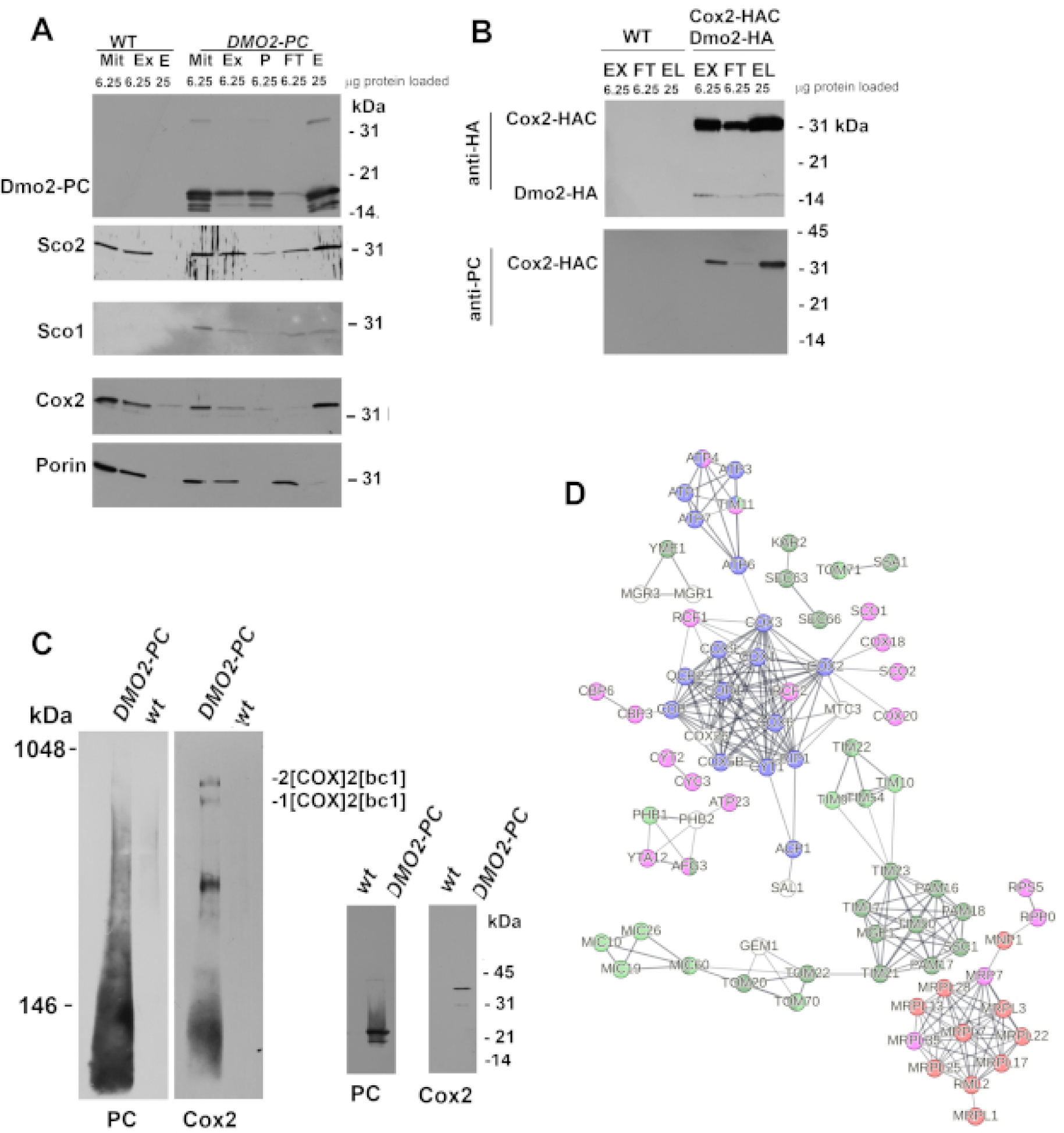
Study of Dmo2p physical interactors. **(A)** Mitochondria (Mit) digitonin extracts from the wild type (WT) and dmo2/DMO2-PC strains were centrifuged, and the pellet (P) saved. The extracts were then incubated with protein C antibody beads, with non-ligand fractions (Ft) and ligands eluted from the beads (E) being collected. The estimated amount of proteins loaded in each lane and the antibodies used in the immunoblots are indicated in panels A and B. **(B)** Mitochondria (Mit) from the wild type (WT) and the strain expressing the *COX2-HAC* allele together with *DMO2-HA* were extracted and analyzed as in (A). **(C)** The pull-down elution from WT and dmo2/DMO2-PC strains was analyzed on a 4-13% non-denaturing acrylamide gel by BN-PAGE (left panel) and on a standard 12.5% acrylamide gel by SDS-PAGE (right panel). The proteins were then transferred to PVDF and nitrocellulose membranes, respectively, and immunoblotted against the indicated antibodies. **(D)** Mass spectrometry analyses and comparisons of the elution fractions obtained from WT and dmo2/DMO2-PC were conducted. The gene ontology map of interactions was obtained using String 12.0 (Szklarczyk et al., 2023); in red: mitochondrial gene expression (GO 0140053); blue: OXPHOS components (KEGG DCE00190); green: protein transmembrane transport and protein quality control (GO 0071806, GO 0006515); yellow: mitochondrial transport (GO 0006839); pink: protein-containing complex assembly (GO 0065003).

Considering the Cox2p-Dmo2p interaction, we investigated whether this interaction occurs solely during the Cox2p maturation process or persists in later stages of COX assembly. To address this, elution fractions from wild-type and Dmo2p-CH mitochondrial extracts, obtained after digitonin extraction, were loaded onto a native 4-13% polyacrylamide gel and immunoblotted with anti-protein C and anti-Cox2 antibodies (Figure 10C). In this panel, *bc*1 and COX supercomplexes with conformations 2[bc1] 2[COX] and 2[bc1] 1[COX] were pulled down in the Dmo2p-CH eluate, suggesting that Dmo2p is transiently associated with these structures. Anti-Cox2p also detected a clear band in the middle of the gel, as well as a smear of bands below 143 kDa; these bands look like COX and Cox2 intermediates (Franco et al., 2018), respectively; however, further experiments are crucial to determine the nature of these bands. Additionally, anti-protein C probing unveiled a broad smear extending from the bottom to the region where the *bc*1 COX supercomplexes were detected, indicating interactions with multiple mitochondrial proteins. Cox2p and Dmo2-PC in the eluates were also detected after SDS-PAGE and immunoblotting.

Finally, we assessed the network of Dmo2p interactors through mass spectrometry analyses of proteins differentially purified from the PC beads in the *DMO2-PC* strain and the wild-type, followed by subsequent enrichment tests. Protein quantification was conducted utilizing the MaxQuant label-free algorithm (LFQ), which employed unique and razor peptides for accurate measurement. A minimum requirement of ≥2 ratio counts was set as essential for valid protein quantification. A total of 188 proteins showed at least a 0.5-fold increment in abundance in the *dmo2/DMO2-PC* transformed mutant compared to the untagged wild-type. Notably, the Ubp16p protein, a deubiquitinating protein of the outer mitochondrial membrane (Kinner and Kölling, 2003), exhibited the highest increment, with a 5.27-fold enrichment. Although this data stemmed from a single experiment, the significant enrichment of Ubp16p supports our hypothesis that Dmo2p may undergo post-translational modifications by ubiquitin (Figure 4A). Additionally, we filtered the gene ontology interactor analyses using a physical subnetwork comprising protein constituents of physical complexes to obtain a refined network of interactions. Therefore, we observed the presence of various constituents along the mitochondrial membranes, including components from mitoribosomes, MIOREX, TIM23 machinery, TOM complex, Micos multi-subunit complex, carrier proteins involved in transport, and protein quality control. Moreover, respiratory chain complex proteins and assembly factors, such as Sco1p, Sco2p, Cox2p, Cox18p, and Cox20p, were detected, suggesting the potential presence of Cox2p assembly intermediates. Structural components of the *bc*1-COX supercomplexes, including Cox1p, Cox2p, Cox3, Cox6, Cox5Ap, Cox9p, Cox26p, Cor1p, Cyt1p, Rip1, Rcf1p, and Rcf2p, were also identified in the eluate, along with assembly factors such as heme lyases Cyt2 and Cyc3, which exhibited significant enrichments of 5-fold and 4.5-fold, respectively. Sal1, known to function alongside Pet9 in ADP/ATP carrier transport, displayed a 2.85-fold enrichment. Interestingly, Pet9, which participates in heme transport and is physically associated with the respiratory supercomplex and the Tim23 machinery, also exhibited enrichment (2.85-fold). In fact, constituents involved in the *bc*1-COX supercomplex, including Rcf2p (1.89-fold enrichment) and Rcf1p (0.69-fold enrichment), were enriched in the Dmo2-CH eluted material. Cox26p, a stoichiometric subunit of the cytochrome *c* oxidase complex required for *bc*1-COX supercomplex formation, was also identified in our mass spectrometry analyses (0.60-fold enrichment). The Dmo2p interactome suggests its involvement in various aspects of mitochondrial metabolism, including potential roles in copper addition to Cox2 through Sco1p and Sco2p, as well as *bc*1-COX supercomplex stabilization.

## Discussion

The present study sheds light on potential investigations to elucidate the redox dynamics and partners of Dmo2p, which may have implications for human health through its homolog DMAC1. Our data support the involvement of Dmo2p in inter-membrane mitochondrial redox metabolism, as evidenced by its requirement in cells subjected to oxidative and heat stresses and the observation of elevated levels of oxidized glutathione in both the mitochondrial inter-membrane space and the cytosol. With its dual localization, Dmo2p may play a role in redox signaling communication between peroxisomes and mitochondria, contributing to cellular responses to oxidative stress and other environmental stimuli.

Furthermore, we have gathered evidence suggesting a potential link between Dmo2p and mitochondrial copper homeostasis. Copper ions are known to be redox-active and highly cytotoxic, needing strict cellular control mechanisms to prevent the accumulation of free intracellular copper. Our findings hint at a possible role for Dmo2p in regulating copper delivery to Cox2p, which could have implications for maintaining cellular redox balance and overall cytochrome *c* oxidase activity.

We observed that overexpression of *DMO2* results in growth impairment in the presence of 12mM CuSO_4_, indicating a potential disruption of copper homeostasis within the mitochondria. Subsequent investigations revealed an interaction between Dmo2p and Sco1p/Sco2p, proteins responsible for distributing copper to Cox2p. Furthermore, the additional copies of *DMO2* partially suppress *cox23* respiratory deficiency, suggesting a potential role for *DMO2* in regulating mitochondrial copper homeostasis and Cox23p function. Cox23p has been implicated in the regulation of mitochondrial copper homeostasis, highlighting its importance in maintaining cellular redox balance (Barros et al., 2004; Longen et al., 2009; Dela Cruz et al., 2016).

In this context, Dmo2p may assist proteins such as Sco1p, Sco2p, Cox2p, and Cox23p in their ability to interact with copper ions or copper-binding partners, thereby modulating mitochondrial copper homeostasis and redox balance. This is consistent with previous findings that the expression of *COX20* and *SCO1* is associated with reduced levels of ROS under oxidative stress conditions, while null mutants display elevated sensitivity to oxidative stress (Keerthiraju et al., 2019; Ekim Kocabey et al., 2019; Rigby et al., 2008).

The complexity of COX copper insertion requires the involvement of multiple proteins, including CX_9_C proteins such as Cox17p, Cox19p, Cox23p, Pet191p, and Coa6p. These proteins accompany Cox2p and Cox1p during copper insertion, thereby preventing uncontrolled ROS generation due to the high reactivity of their intermediates (Glerum et al., 1996; Nobrega et al., 2002; Longen et al., 2009; Nývltová et al., 2022; Veniamin et al., 2011; Khalimonchuk et al., 2007; Geldon et al., 2021). The CX_2_C motif present in *DMO2* homologs is crucial for Dmo2p function, likely playing a role in coordinating protein interactions or redox processes. This motif is also found in DMAC1 (TMEM261), which has been implicated in the assembly of respiratory complex I distal arms and supercomplex formation (Stroud et al., 2016).

In our study, we observed that Dmo2p interacts with the yeast *bc*1-COX supercomplexes and several components of the mitochondrial intermembrane transport system, suggesting its involvement in mitochondrial function and biogenesis. These interactions were identified through a combination of techniques, including sedimentation gradients, co-immunoprecipitation and mass spectrometry analysis.

Recent quantitative mapping of mitochondrial protein assemblies, termed MitCOM (Schulte et al., 2023), revealed that Dmo2p’s top correlating protein profile includes Cbp4p, Cyt2p, Sco1p, Sco2p, and Cox20p, which were also identified in our mass spectrometry analysis of Dmo2p co-immunoprecipitated material. Cbp4p and Cyt2p are known to participate in the biogenesis of the cytochrome *bc*1 complex assembly. Cbp4p acts as an assembly factor that associates with newly synthesized cytochrome *b* (Gruschke et al., 2011), while Cyt2p serves as the cytochrome *c*1 heme lyase, linking heme covalently to apocytochrome *c*1 (Zollner et al., 1992).

These findings support the hypothesis that the Dmo2p may have multiple clients, potentially including proteins involved in transporting metal ions, heme, and components of the TIM23 machinery through the mitochondrial inner membrane; the identification of Cox2p and its partners in our study further strengthens this hypothesis, highlighting the diverse roles of Dmo2p in mitochondrial respiratory capacity.

## Material and Methods

### Yeast strains and growth media

The genotypes and sources of yeast strains used in this study are listed in Table 1. Yeast strains were maintained in YPD (1% yeast extract, 2% peptone, 2% glucose), YPGal (1% yeast extract, 2% peptone, 2% galactose), YPEG (1% yeast extract, 2% peptone, 2% glycerol, 2% ethanol), minimal glucose (2% glucose, 0.67% yeast nitrogen base (YNB) without amino acids, supplemented with auxotrophic requirements), YNO (0.1% oleic acid, 0.1% Tween 20, 0.67% complete YNB, 0.5% yeast extract), and sporulation media (0.5% yeast extract, 1% peptone, 0.05% glucose); all reagents, unless otherwise indicated, were obtained from Sigma-Aldrich (Saint Louis, MO).

**Table 1.**
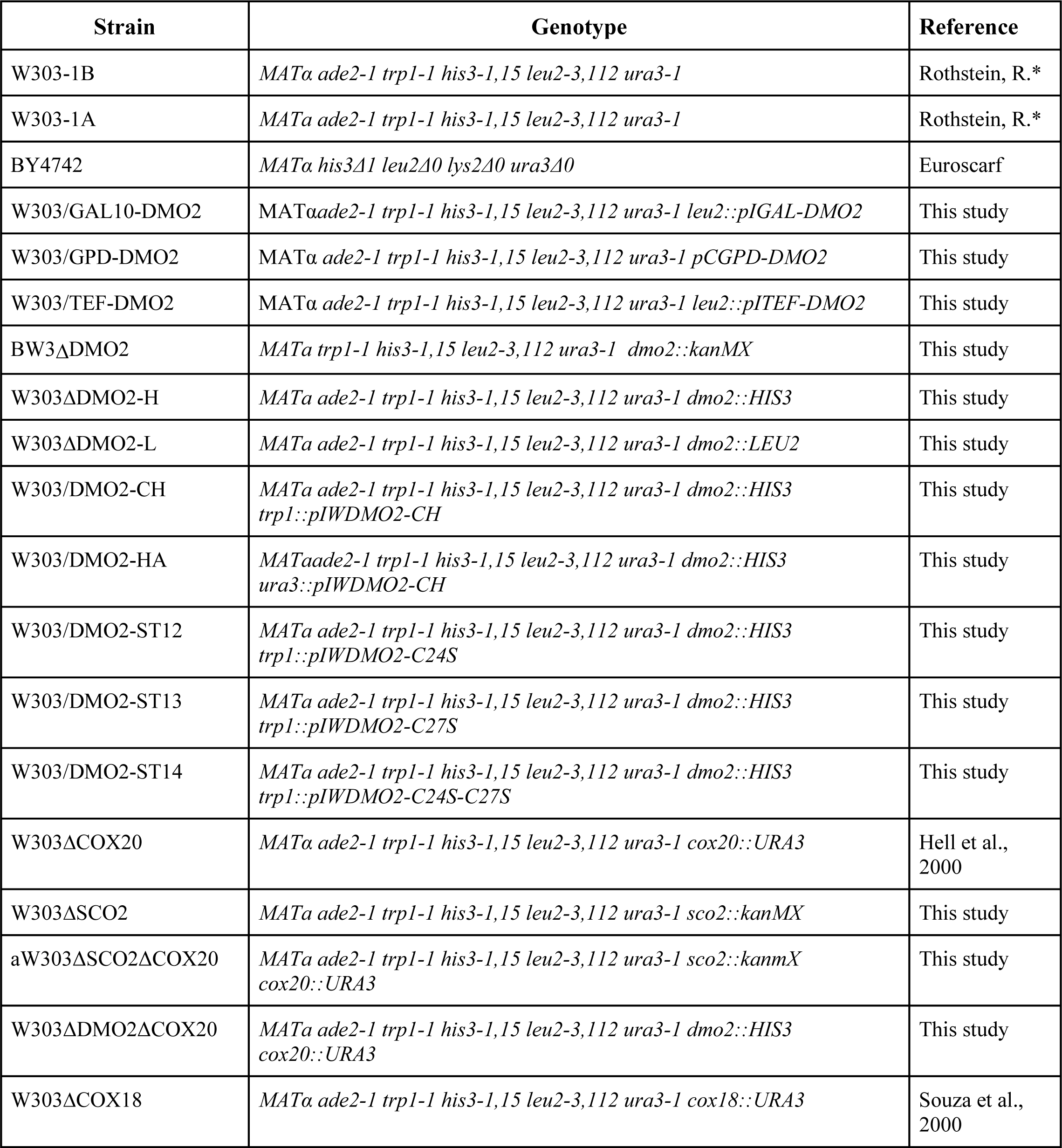
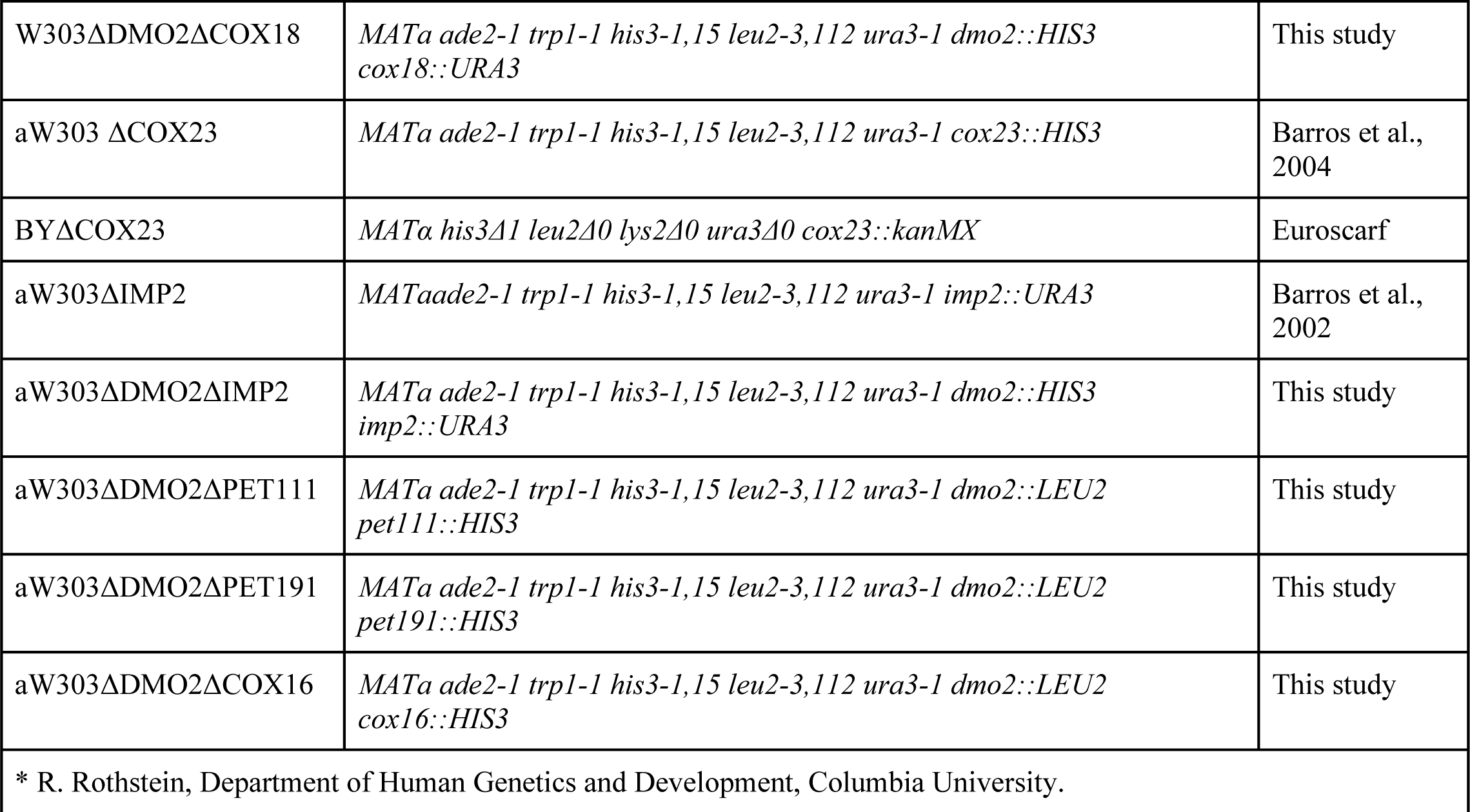
Yeast strains and relevant genotype used in this work.

### Construction of DMO2 alleles

The *DMO2* wild-type allele was amplified from nuclear DNA with primers DMO2-F0 (5’ GGCCTGCAGACTCACCCCTTAGCCT) and DMO2-2 (GGCGGATCCATTGAATGCATGCGATAAC); the 1110 bp PCR product was double digested with BamH1 and Pst1 and cloned in pUC18, YIp352, YEp351 (Hill et al., 1986), all previously cut with the same enzymes yielding pUC18/DMO2, pDMO2/ST1 and DMO2/ST2. The pUC18/DMO2 was used as a template for a new PCR cycle with the bi-directional primers GGCAGATCTTGTTTTTCACCACTTGCT and GGCGGATCCATTGAATGCATGCGATAAC, resulting in a clean deletion of *DMO2*. The linear product containing the *DMO2* flanking region in pUC18 was digested with Bgl2 and ligated to *HIS3* and *LEU2* on approximately a 1.2 kb BamH1 fragment of DNA. The *dmo2::HIS3* and *dmo2::LEU2* null alleles were recovered from the recombinant plasmids as a 1.5 kb Pst1-BamH1 fragment and were substituted by one-step gene replacement for the wild-type allele in the yeast strain W303-1A.

*DMO2* C-terminal was fused to the HA tag by PCR amplification with DMO2-F0 and 5’GGCATGCATTCAAGCGTAGTCTGGGACGTCGTATGGGTAGTTAGCCTGTGTTTCACCATC (DMO2-HA) primers. This PCR product contains the *DMO2* endogenous promoter, and it was cloned into YIp352 and afterward transferred to YIp349 (Hill et al., 1986). A second PCR amplification using DMO2-HA and 5’ GGCGGATCCATGAGTAATATTTTGGCAGT primers amplified only the *DMO2* coding sequence. This PCR product was cloned into YIp351-TEF1, YIp351-GPD, and YIp351-GAL (Zampol et al., 2010) in which the coding sequence of *DMO2-HA* allele is under control of the *GAL10* promoter, the translation elongation factor 1α promoter (*TEF1*) and the glyceraldehyde-3-phosphate dehydrogenase promoter (*GPD*) (Mumberg et al., 1995). The DMO2-PC fusion was obtained using primers DMO2-F0 and 5’ GGCCTGCAGGTTAGCCTGTGTTTCACCATC primers, digested with Pst1 and cloned in frame with the protein C sequence present in YIp351-CH plasmid.

The cysteine point mutants were created by site-directed mutagenesis using two PCR reactions, in both reactions the primer is annealed to the desired region, generating the mutation and an Nco1 site. For the first reaction, we combined a pair of primers to generate the mutation; therefore for the allele *dmo2C24S,* we used DMO2-2 with 5’ GGCCCATGGATTCCGTTCCTTGTCAAGTC; to generate *dmo2C27S:* DMO2-2 with 5’ GGCCCATGGATTGCGTTCCTTCTCAAGTC; to generate *dmo2C24S C27S:* DMO2-2 with 5’ GGCCCATGGATTCCGTTCCTTCTCAAGTC. The second reaction was the same for the three mutants and used the pair DMO2-F0 with 5’GGCCCATGGTCTCTTCTTTTTCCAGCTCC. We digested the products of the first reaction with Nco1 and BamH1 and the products of reaction 2 with Nco1 and Pst1. The double-digested fragments were cloned into YIp352, YIp349, and YEp351, previously digested with BamH1 and Pst1 (Hill et al., 1986).

### Mitochondrial Protein Synthesis

Yeast cell cultures were cultured in rich galactose media. Mitochondrial translation products were labeled in whole cells with a mixture of ^35^Smethionine/cysteine (7 mCi/mmol) in the presence of cycloheximide. Cells were lysed and protein extracted as previously described (Santos et al., 2021). Total cellular proteins were separated by SDS-PAGE on 17.5% polyacrylamide or 6M UREA-PAGE on 12.5% polyacrylamide gel systems.

### Antibodies

We acknowledge the donation of the antibodies used in this work: A. Tzagoloff – Columbia University (Anti-Rip1, Anti-β subunit of F1, Anti-Cox1, Anti-Cox2, Anti-Cytb2, Anti-Kgd2, Anti-Sco1, Anti-Sco2); JP D-iRago – Univ. Bordeaux (Anti-Atp6); C. Clark – UCLA (Anti-Coq4) W. Girzalsky - Ruhr University (Anti-Fox3); M. Demasi – I.Butantã (Anti-Prx1); Martin Ott - Stockholm University (Anti-uL24). Anti-porin was bought from Thermofisher (Waltham, MA - USA) and anti-Cox2 from Abcam (Cambridge, MA – USA). Anti-ubiquitin P4D1 (sc-8017) from Santa Cruz Biotechnology (Dallas, TX – USA)

### Mitochondria isolation and fractionation

Yeast mitochondria were prepared by the method of Herrmann et al. (1994) using Zymolyase 20T – Zymo-Research (Irvine, CA) to obtain spheroplasts. Protein concentrations were determined by the method of Lowry et al (1951). Six mg of mitochondrial preparation were solubilized in 400 µL of an extraction buffer (20 mM Hepes pH 7.4, 25 mM KCl, 0.5 mM PMSF, 1% Dodecyl maltoside X100 (or 2% Digitonin), 5 mM MgCl_2_) and were centrifuged at 27,000 × g for 15 minutes and layered onto a 7 to 20% linear sucrose gradient containing 10 mM Tris-Cl pH 7.5, 0.5 mM EDTA and 0.05% Triton X-100. The gradient was centrifuged at 54,000 rpm for 5 hours in a Beckmann SW55Ti rotor. The gradient was collected by gravity in 15 equal fractions and each fraction was assayed for hemoglobin by absorption at 410 nm and for lactate dehydrogenase by measuring NADH-dependent conversion of pyruvate to lactate. The organelle fraction from oleate-grown cells (about 1.4 mg of protein) was analysed in a discontinuous sucrose gradient (cushion of 0.4 ml of 60% sucrose, 1.05 ml each of 50%, 46%, 43%, 40% and 0.4 ml of 35% sucrose; McCammon et al., 1990). The gradient was centrifuged at 100,000 x g for 6 hours.

### Enzymatic assays of mitochondrial respiratory complexes

NADH-cytochrome *c* reductase activity was measured by following the increase in absorbance at 550 nm due to the reduction of cytochrome *c* in the presence of 10mM NADH (NCCR). COX was assayed by following the oxidation of ferricytochrome at 550nM (Tzagoloff et al., 1975). ATPase in-gel assays were performed as described elsewhere (Franco et al., 2020). 50μg of mitochondria were extracted with 2% digitonin and loaded onto a 4–13% non-denaturing blue native gel (BN-PAGE). O_2_ consumption assay was monitored over time using a computer interfaced Clark electrode operating at 25°C with continuous stirring in the presence of 0.5mM NADH and 0.1mg of mitochondria extract.

### Mitochondrial matrix glutathione oxidation state measurements

W303-1B and *dmo2*Δ mutants were transformed with the plasmid p416-Su9-Grx1, which addresses to the mitochondrial matrix the modified ro-GFP (Morgan et al., 2011). We follow a protocol described elsewhere (Calabrese et al., 2019). The transformants were grown in galactose minimal media without uracil for plasmid selection at 30°C. The cells were harvested by centrifugation at 1,500 × g for 3 minutes at room temperature and subsequently suspended in 0.1 M sorbitol, 0.1 M Tris-HCl pH 7.4, 0.1 M NaCl to a final concentration of 1.5 OD600 units/ml; 180 μl of cells were transferred to appropriate microplates. To one well, diamide was added as the fully oxidized control. To a second, DTT was added as the fully reduced. Fluorescence was recorded using filter optics at excitation wavelengths of 410 ± 5 nm and 482 ± 8 nm, and an emission wavelength of 530 ± 20 nm. A basal response was measured for 10 minutes, then the sample cell wells were challenged with H_2_O_2_ 1mM and the response was followed for 60 minutes.

### Mass spectrometry and Bioinformatics analysis

Mass spectrometry analyses were performed at the Redox Proteomics Core of the Mass Spectrometry Resource at Chemistry Institute, University of São Paulo; the results were processed using the Perseus software suite (http://www.perseus-framework.org/). Data were transformed [log2(x)] and filtered so that for each protein, at least one group (Dmo2-PC/wild-type) contained a minimum of 50% valid values. Subsequent steps in Perseus included a two-sample t-test to compare the means of the label-free quantitation (LFQ) intensity values between the Dmo2-PC and wild-type groups.

### Miscellaneous procedures

Protein concentrations were determined by the method of Lowry (Lowry et al., 1951), and were suspended with the Laemmli buffer (Laemmli, 1970) for SDS-PAGE blot analyses. For BN-PAGE, mitochondrial proteins were extracted with 2% final concentration of digitonin and separated on a 4-13% polyacrylamide gel under non-denaturing conditions; proteins were transferred to a PVDF membrane and probed with antibodies indicated in the Figures (Franco et al., 2019). The antibody-antigen complexes were visualized with coumaric acid, luminol and hydrogen peroxide mixture. Densitometry analysis of the bands was done using the histogram component of GNU Image Manipulation Program GIMP 2.10.28. GraphPad Prism software was used for graphs and data analyses by one-way multiple pairwise Tukey tests.

## Acknowledgments

We would like to acknowledge Professor Luis Netto (IB-USP) for providing reagents and granting us access to the fluorescence reader equipment. Dr. Marilene Demasi for the discussions ascertains our results. We thank Professors Paolo Di Mascio and Graziella E. Ronsein for the Mass spectrometry analyses performed at the Redox Proteomics Core of the Mass Spectrometry Resource at Chemistry Institute, University of São Paulo (FAPESP grant numbers 2012/12663-1, 2016/00696-3, CEPID Redoxoma 2013/07937-8), and Dr. Mariana P. Massafera for her technical assistance. This work was supported by grants and fellowships from Fundação de Amparo à Pesquisa de São Paulo (FAPESP 2020/05812-7, 2013/07937-8), Conselho Nacional de Desenvolvimento Científico e Tecnológico (CNPq 305054/2022-8), and Coordenação de Aperfeiçoamento de Pessoal de Nível Superior - Brasil (CAPES) - Finance Code 001. Maria Kfouri, Jhulia Chagas, and Letícia Franco are fellowship recipients from FAPESP (FAPESP-2022/02744-6, 2022/08559-6, 2019/02799-2).

## Author Contributions

MAKMS – Performed the experiments, analyzed the data, and prepared the digital images.

LVRF - Conceived and designed experiments.

JACH - Performed the experiments.

FG - Performed the experiments, analyzed the data.

MHB – Conceived, designed and performed experiments; prepared the digital images and drafted the manuscript.

