## supplemental file for "*Saccharomyces cerevisiae* Dmo2p is involved in the mitochondrial copper metabolism and is required for the stability of newly translated Cox2p"

**Figure S1** – *DMO2* proteomics from y3kproject.org website. Screenshots of protein abundance in volcano plots from yeast cultures grown on conditions that favored the respiratory metabolism A) *dmo2Δ* mutant proteome protein abundance in comparison to the wild-type strain, B) Dmo2p (Ydl157c) distribution in the knockout Euroscarf collection. C) Correlation between protein abundance in *mrpl11Δ* and *dmo2Δ* mutants, D) Correlation between protein abundance in *mef2Δ* and *dmo2Δ* mutants.

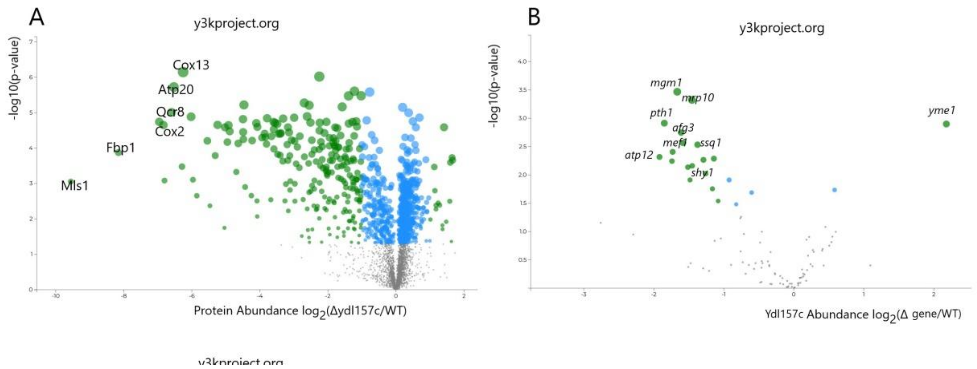


**Figure S2** – Global genetic interaction of *dmo2Δ* mutant obtained from thecellmap.org website. Screenshots of *dmo2Δ* profile similarity (A) and negative genetic interactions (B). Dots painted in green represent mitochondrial protein mutants.

**A** **B**

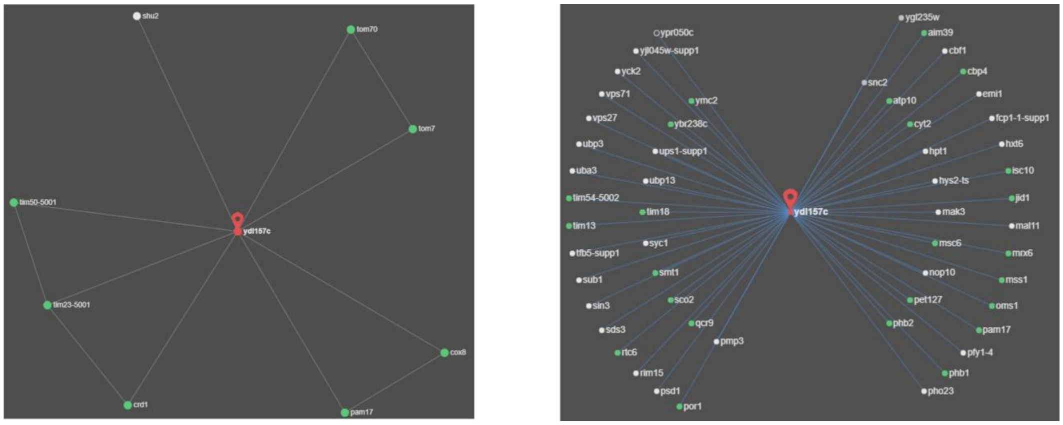

**Figure S3** – Dmo2p sequence alignments, and modeling. (A) AlphaFold protein structures modeled for human DMAC1 and Dmo2p, the conserved cysteine are indicated (< Cys) as well as the N and C ends of each protein. (B) Clustal Omega alignment of human DMAC1 with Dmo2p (Ydl157c) and *Schizosaccharomyces pompe* Tam6p. (C) Dmo2p conservation residues map along the fungi obtained using the EV-coupling server.

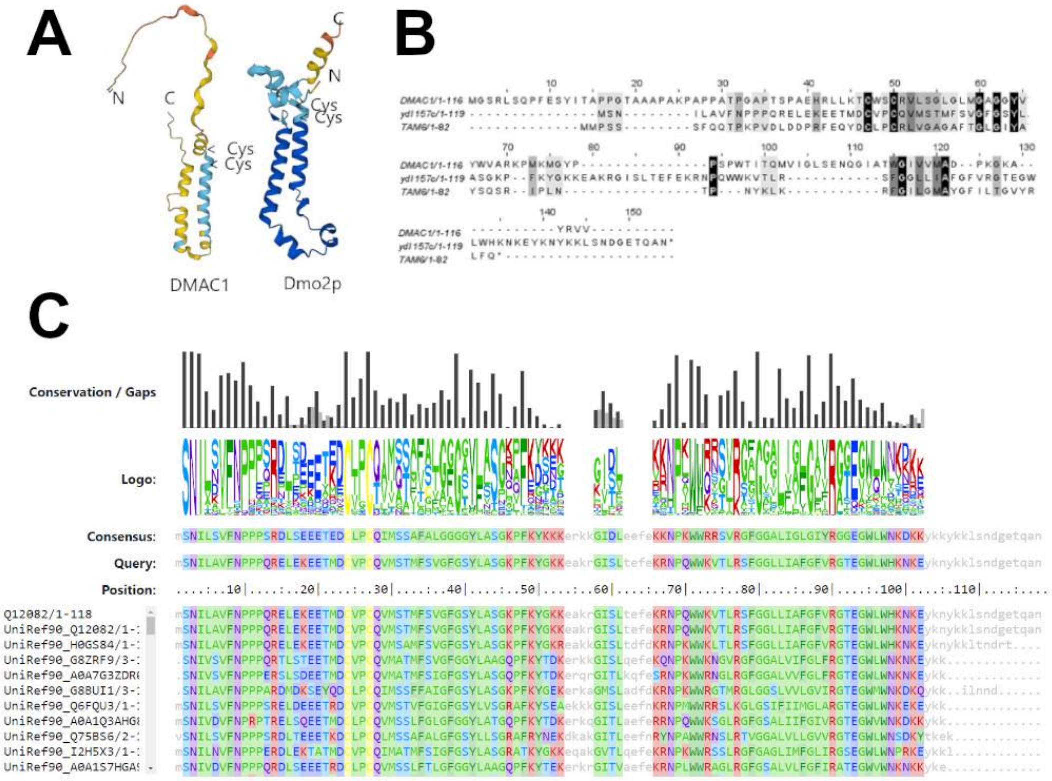

**Figure S4**:. **(A)** Hypochlorite, hydrogen peroxide, $ZnCl_2$, $CuSO_4$ and $FeCl_3$ stress challenges. Cells were incubated for 5 hours in different concentrations of hypochlorite (left panel) and for 2 hours at different temperatures (right panel) before being plated in the indicated medium and grown at 30ºC for 2 days. Analyzed strains were parental W303-1A, *DMO2* mutants *dmo2::LEU2* (dmo2ΔL) and *dmo2::HIS3* (dmo2ΔH), and W303 overexpressing *DMO2* with *GPD* promoter (WT+DMO2). **(B)** Serial dilution test in rich glucose medium (YPD), rich galactose medium (YPGAL) and rich ethanol-glycerol medium (EG) with strains W303-1A and overexpressing *DMO2* in *GAL10* promoter (W303+pIGAL-DMO2) and in *GPD* promoter (W303+pCGPD-DMO2). Plates were photographed after 1 day of growth.

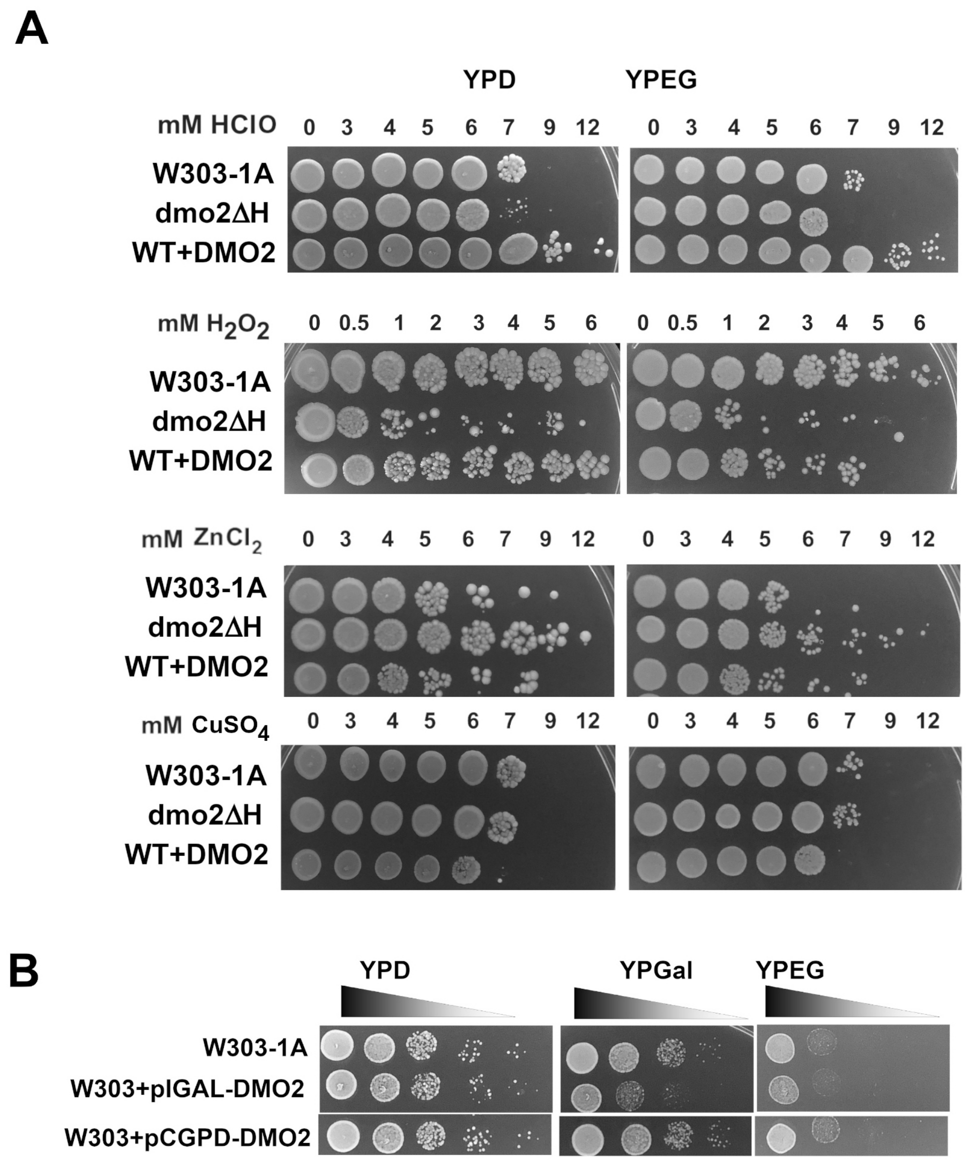

**Figure S5** – Heat shock response comparison. Equal amounts of cells at logarithmic growth from the wild-type culture and *dmo2* null mutant strain (ydl157c∆) were incubated for 15 minutes at room temperature (RT) and at 38°C. Afterwards, the cells were maintained at 47°C for two hours; the survival rate was scored by dividing the number of colonies units by the estimated number of cells spread on the plate.

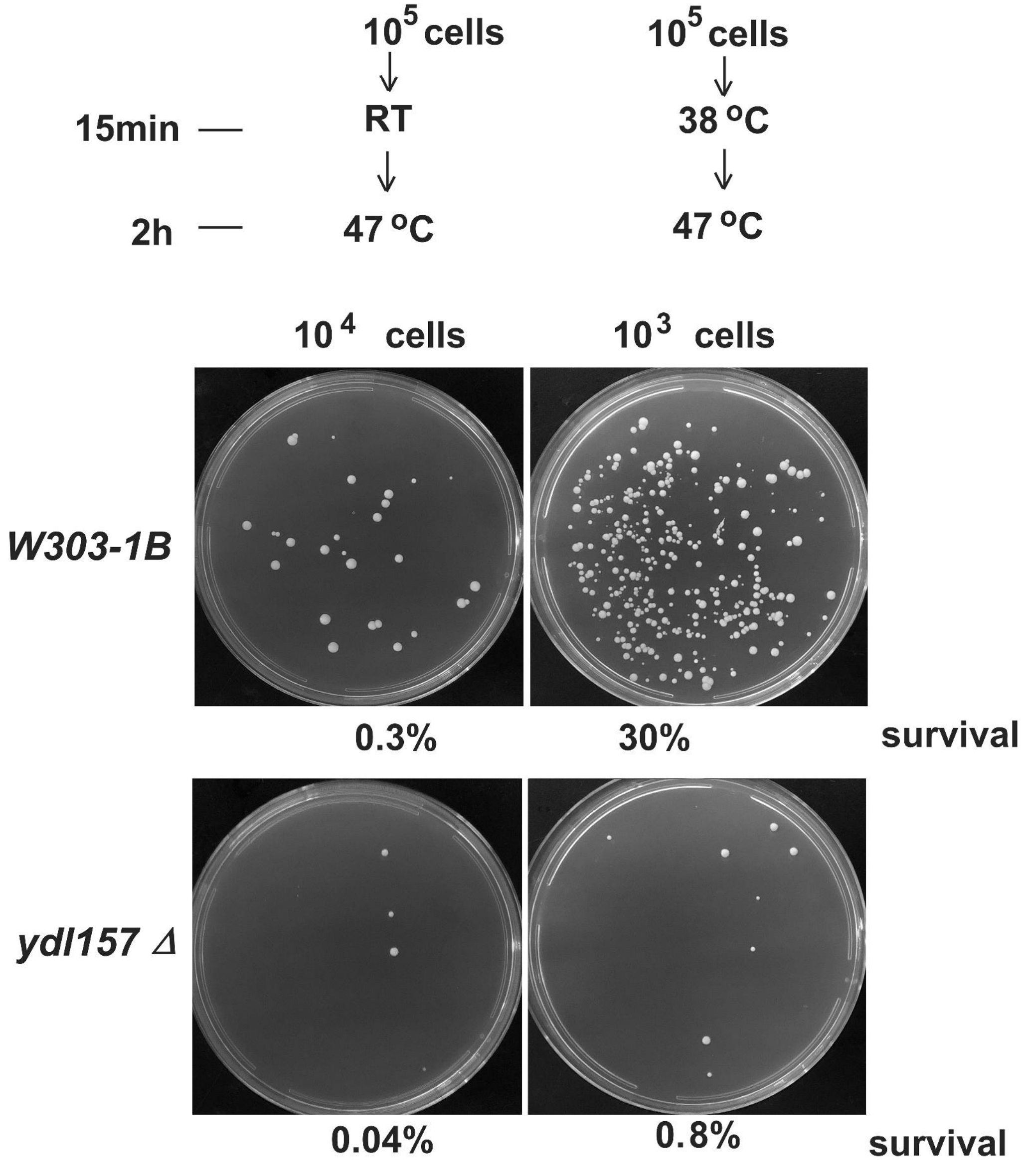

**Figure S6** – Growth properties of the *cox20* mutant strains. The wild-type (WT) and the indicated mutants were spotted on rich glucose media (YPD) and rich non-fermentable ethanol-glycerol media (YPEG). YPD plates were photographed after two days, and YPEG plates after 2 or 7 days as indicated.

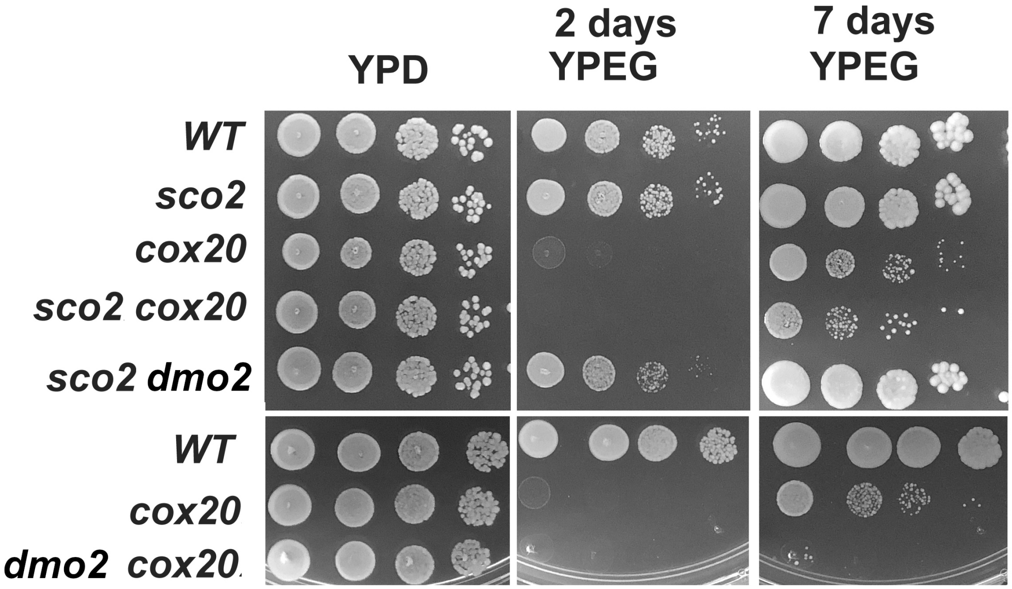

**Figure S7** 50μg of WT and *dmo2* mitochondria isolated from cells grown at the indicated temperatures were extracted with 2% digitonin. Extracts were loaded to a non-denaturing 4-13% acrylamide gradient BN-PAGE gel. Membranes were probed with antibodies anti-Cox1, anti-Cox2, anti-Cor1 and anti-Cob respectively.

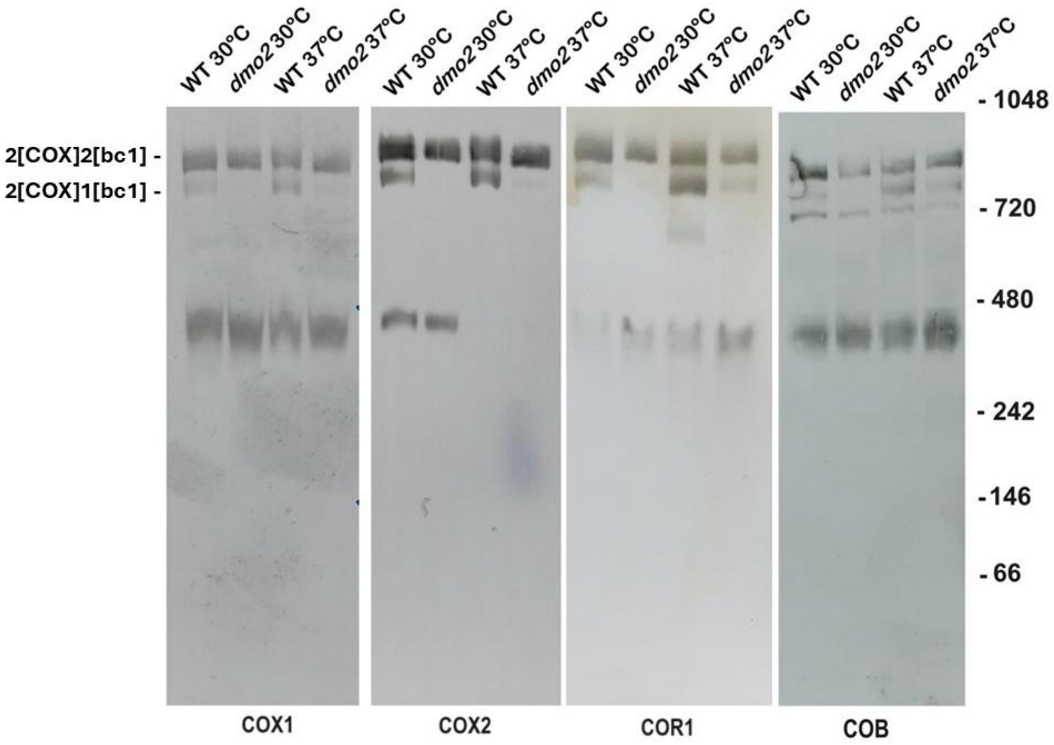

**Figure S8 –** (A) Steady state protein levels; Rip1 and Cor1 are components of respiratory complex III (CIII); Cox1 and Cox2 are components of respiratory complex IV (CIV); Atp6 is a component of ATP synthase, or complex V (CV); uL29 is a component of the mitoribosome large subunit; Prx1 is a soluble mitochondrial protein and Porin is a protein from the mitochondrial outer membrane. (B) Labeling of mitochondrially encoded proteins as described in Figure 6, for the wild type strain (WT) *yme1* null mutant, *dmo2 yme1* double mutant, and *dmo2* mutant.

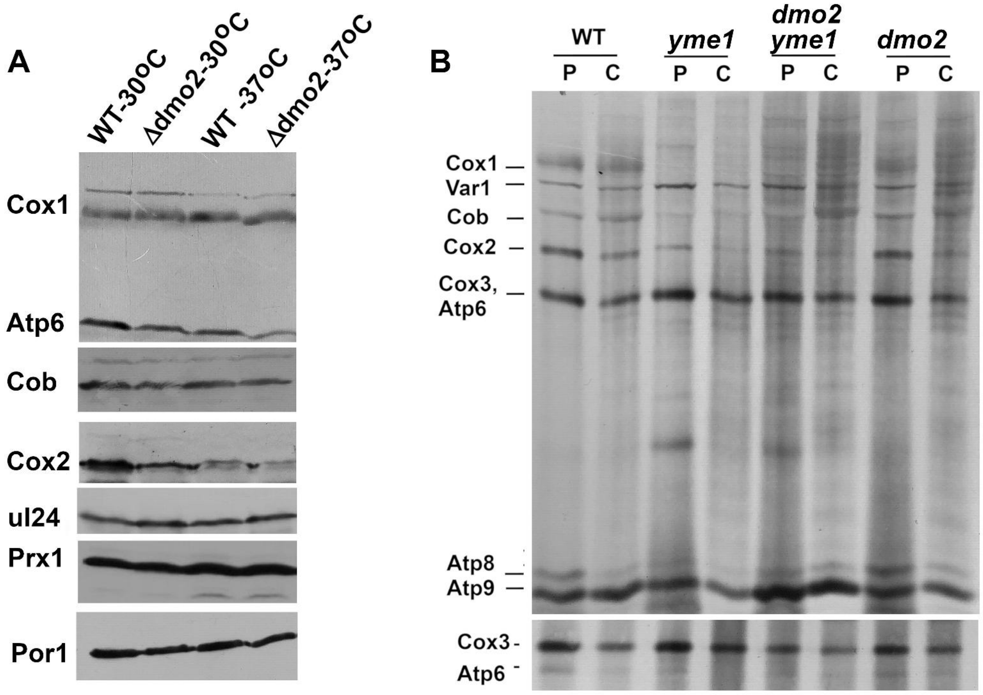

**Figure S9** – *DMO2* suppression is restricted to *cox23* mutants. DMO2 overexpression in multicopy plasmids (YEp351-DMO2) and GPD-DMO2 constructs were tested in mutant members of CyC family *cox17*, *cox19* and *cox23*

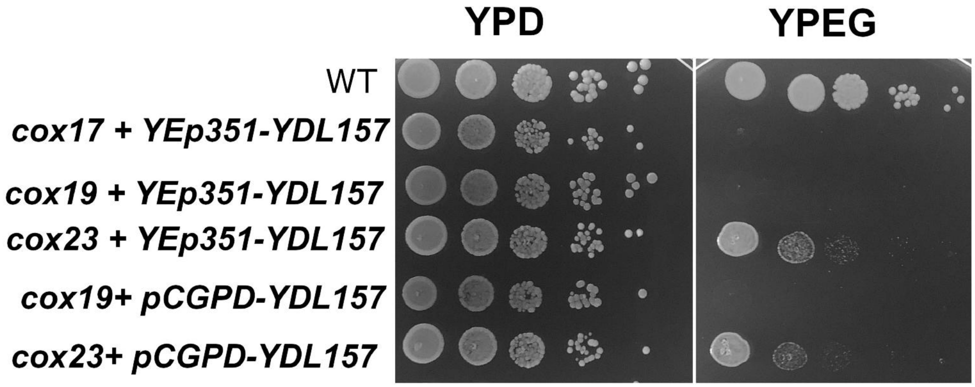
